# Intrinsic and Rho-dependent termination cooperate for efficient transcription termination at 3’ untranslated regions

**DOI:** 10.1101/2022.07.21.500918

**Authors:** Ezaz Ahmad, Varsha Mahapatra, Vanishree V M, Valakunja Nagaraja

## Abstract

The intrinsic, and the Rho-dependent mechanisms of transcription termination are conserved in bacteria. Generally, the two mechanisms have been illustrated as two independent pathways occurring in the 3’ ends of different genes with contrasting requirements to halt RNA synthesis. However, a majority of the intrinsic terminators terminate transcription inefficiently leading to transcriptional read-through. The unwanted transcription in the downstream region beyond the terminator would have undesired consequences. To prevent such transcriptional read-through, bacteria must have evolved ways to terminate transcription more efficiently at or near the termination sites. We describe the participation of both the mechanisms, where intrinsic terminator and Rho factor contribute to prevent transcriptional read-through. Contribution from both the termination processes is demonstrated at the downstream regions of the genes both *in vitro* and *in vivo* in mycobacteria. Distinct patterns of cooperation between the two modes of termination were observed at the 3’ untranslated regions of the genes to ensure efficient termination. We demonstrate similar mode of operation between the two termination processes in *Escherichia coli* suggesting a likely prevalence of this cooperation across bacteria. The reporter system developed to assess the Rho – intrinsic termination collaboration *in vivo* for mycobacteria and *E. coli* can readily be applied to other bacteria.

## Introduction

Accurate and efficient transcription termination is an important step to generate functional RNA transcripts. The two distinct transcription termination mechanisms in bacteria are intrinsic and Rho-dependent termination (RDT). Intrinsic termination, often termed as simple termination or factor-independent termination is guided by the sequences that form a G/C-rich hairpin stem followed by a U-tract, which stimulates transcription elongation complex (TEC) to pause and dissociate (1, 2). Rho-dependent transcription termination involves the well-conserved termination factor Rho that halts RNA synthesis downstream of the coding sequences to prevent unnecessary transcription read-through (3–5). While intrinsic termination is conserved in all bacterial species, Rho-dependent transcription termination is found in a majority of the bacteria. However, the factor (Rho) seems to be essential for survival only in some bacterial species possibly indicating a major role played by the factor-dependent termination in these species (6–11). In organisms where termination has been studied, there seems to be a preference either for intrinsic termination or RDT as a predominant mode of transcription termination (12–14). In *E. coli*, intrinsic termination appears to be the major mode; and about 20-30% of its genes are terminated by Rho (15, 16). Nevertheless, Rho is found to be an essential gene in *E. coli* (17). In contrast, Rho is not essential in *Bacillus subtilis* and the organism seems to be overly reliant on intrinsic termination (13, 18, 19). In mycobacteria, intrinsic terminators are underrepresented and Rho is essential (10, 14, 20, 21), suggesting their dependence on RDT.

Efficient transcription termination at the end of a gene is important for the regulation of gene expression. Barring intrinsic terminators of rRNA and a few others, a majority of the genes terminate transcription inefficiently. Their termination efficiencies range from as low as 14% to 70% (12, 22–25). This inefficiency appears to arise from the weaker terminator structure. A canonical intrinsic terminator consists of a strong G/C-rich stem followed by 7 to 9 U’s whereas a non-canonical intrinsic terminator is comprised of fewer number of U residues and secondary structures with higher ΔG(20, 26–30). The inefficiency of intrinsic terminators is exacerbated in mycobacteria. As such, mycobacteria suffer from the paucity of intrinsic terminators which is further compounded by the lack of canonical terminators (12, 20). Inefficient transcription termination and read-through can have deleterious consequences to cells due to undesired or untimely transcription of downstream genes (10, 25, 31). Hence organisms have evolved mechanisms to bring about more efficient termination *in vivo* when intrinsic terminators are by themselves inefficient. Our previous study suggested suppression of transcription read-through *in vivo* despite the presence of weak intrinsic terminators in mycobacteria, hinting at the role of additional players in termination (23).

In this study, we have explored the possibility of the two different modes of termination coming together for better control of transcription termination. We demonstrate that both intrinsic and Rho-mediated termination processes cooperate in mycobacteria as well as in *E. coli* to ensure more efficient transcription termination.

## Results

### Rho contributes to transcription termination at intrinsic terminators

Rho controls transcriptional read-through in bacteria (10, 15). However, the participation of RDT at an intrinsic terminator or preceding it to prevent transcriptional read-through at the 3’ untranslated region (UTR) of genes has not yet been addressed. In a previous study, how 3’ end processing of mRNA masked the RDT downstream of the intrinsic terminators was illustrated by Wang et al (32), indicating the participation of RDT downstream of intrinsic termination site, possibly at varied distance (32). We surmised that a combined action of the two processes at a site is of importance especially when intrinsic terminators are inefficient by themselves to cause termination. First, we estimated the contribution of intrinsic terminator and Rho in termination by using a mRuby reporter fused downstream of an intrinsic terminator in a vector transformed in a Rho-conditional knockdown (CKD) strain as described in Materials and methods (Figure 1A). In this vector transcription of the reporter is driven by hsp60 promoter and rrnBT2 at the end serves to terminate transcription (see Materials and methods). For the analysis we employed non-canonical intrinsic terminators from *Mycobacterium tuberculosis* (Mtb) which have fewer number of U residues following the weaker stem-loop structure (higher ΔG) than canonical intrinsic terminators (Supplementary Table S1) (23). To investigate Rho’s contribution to termination with these terminators, the *Mycobacterium smegmatis* (Msm) Rho-CKD strain was constructed (Supplementary Figure 1) in which Rho expression levels were reduced (see Materials and methods). In this *dcas9* based system, depleting the levels of Rho by increasing aTc concentration caused growth retardation and arrest (Supplementary Figure 1B), confirming the essentiality of the gene in mycobacteria (10). In the Rho depleted strain high levels of mRuby expression is observed in absence of intrinsic terminator (Figure 1B, C, D, and Supplementary Figure 2A). However, in presence of intrinsic terminator upstream of the reporter gene, the expression was significantly reduced indicating that intrinsic termination is decreasing mRuby expression (Figure 1B, C, D, and Supplementary Figure 2A) reducing the transcription read-through. The possibility that plasmid copy number is altered in Rho depleted cells contributing to the alteration in mRuby levels was examined by estimating hygromycin resistant gene (Hyg^R^) transcripts in control and Rho depleted strain. As Hyg^R^ transcripts remained unaltered in these experimental conditions, the plasmid copy number is not affected in Rho-CKD strain indicating that the decrease in mRuby expression is due to the introduction of the terminator (Supplementary Figure 2B). Employing this reporter system, we measured the termination efficiency contributed by both intrinsic terminator and Rho at the 3’ UTR of *tuf, cyp144*, and *bfrB* genes of Mtb. The difference in the percentage of mRuby expression in the presence and absence of intrinsic terminator in the normal and reduced level of Rho indicate the individual contribution of both the modes of termination (Supplementary Figure 2C). For *tuf* intrinsic terminator, the termination efficiency was estimated to be 78 ± 2.7 %. However, at depleted Rho levels, the efficiency reduced to 52.4 ± 3.9 % (Figure 1E), indicating the contribution of Rho in the termination of *tuf* gene. Rho-contributed increase in total transcription termination was also observed for *cyp144* and *bfrB* genes as depicted in the table (Figure 2 and Supplementary Figure 3). In contrast, *rrnBT1* which harbours a strong intrinsic terminator (24) did not show any change in termination efficiency in the Rho knockdown background indicating that Rho has rather an insignificant role in transcription termination at strong terminators (Figure 2 and Supplementary Figure 3).

**Figure 1.**
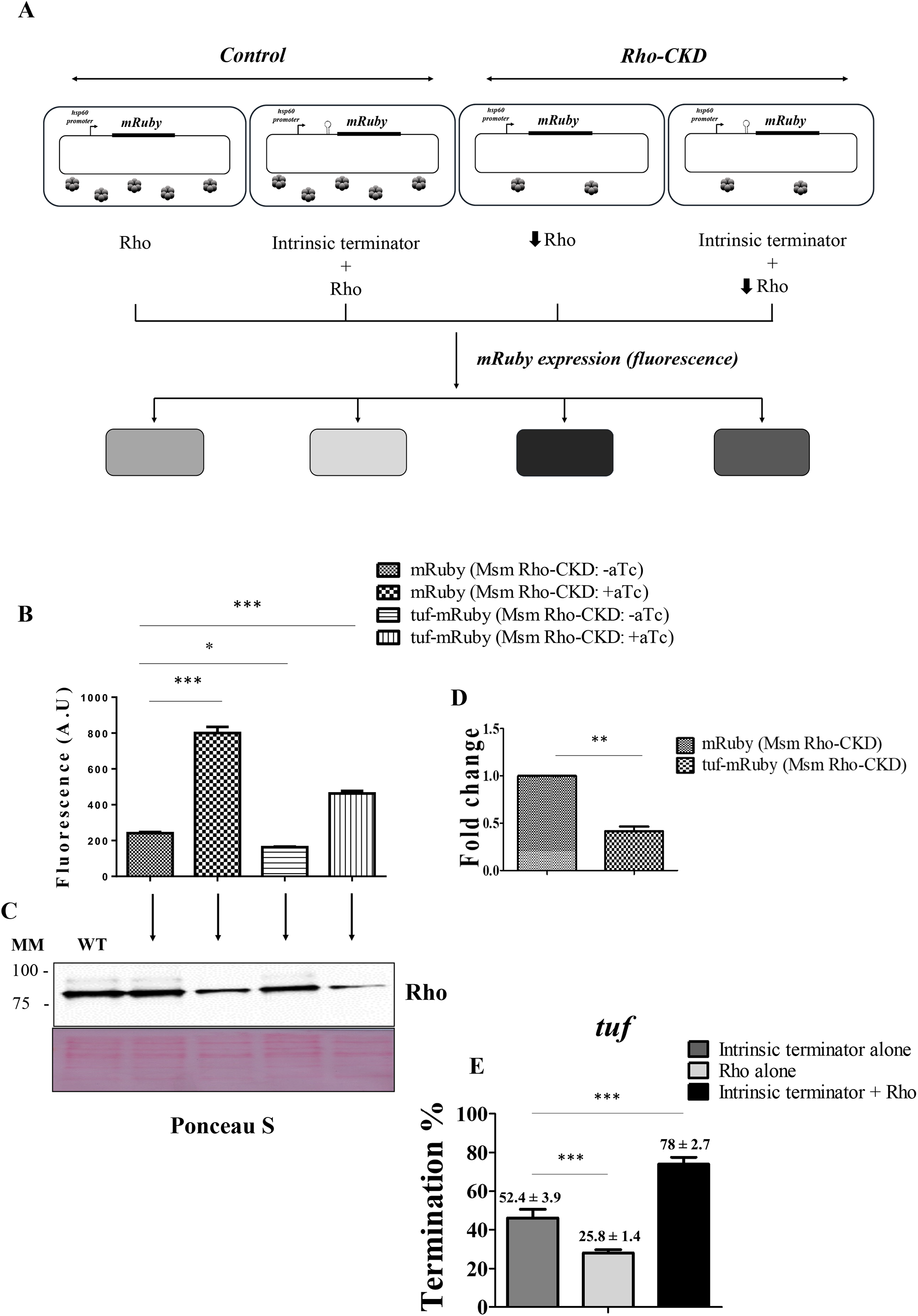
Reporter assay to determine the contribution of intrinsic and RDT. **(A)** Schematic representation of experimental strategy to estimate intrinsic and Rho-dependent termination (RDT) using mRuby fluorescence as a reporter. The reduction in mRuby fluorescence is dependent on the presence of an intrinsic terminator and Rho levels in the cells. Dark to light shade in the schematic represents the intensity of mRuby expression in bacteria in the given conditions. **↓** represents Rho depletion. **(B)** Comparison of mRuby expression level in the presence and absence of *tuf* intrinsic terminator in Msm Rho-CKD strain at normal and depleted levels of Rho. **(C and D)** Immunoblotting and qRT-PCR representing Rho protein and mRuby RNA levels respectively in these experimental conditions. MM represents molecular marker (kDa) and Ponceau S staining is shown as a loading control. **(E)** Contribution of intrinsic terminator alone, Rho alone, and intrinsic terminator + Rho in transcription termination of *tuf* was estimated as a percentage of mRuby fluorescence from transcriptional read-through in Rho-CKD background, in the presence and absence of intrinsic terminator (mRuby Msm Rho-CKD +aTc) – (*tuf*-mRuby Msm Rho-CKD +aTc or – aTc) / (mRuby Msm Rho-CKD +aTc). The data represented is mean ± standard deviation (SD) from the three independent experiments. The statistics were calculated using one-way ANOVA followed by Tukey multiple-comparisons test, *p<0.05, **p<0.001,***p<0.0001, ‘ns’ stands for not significant.

**Figure 2.**
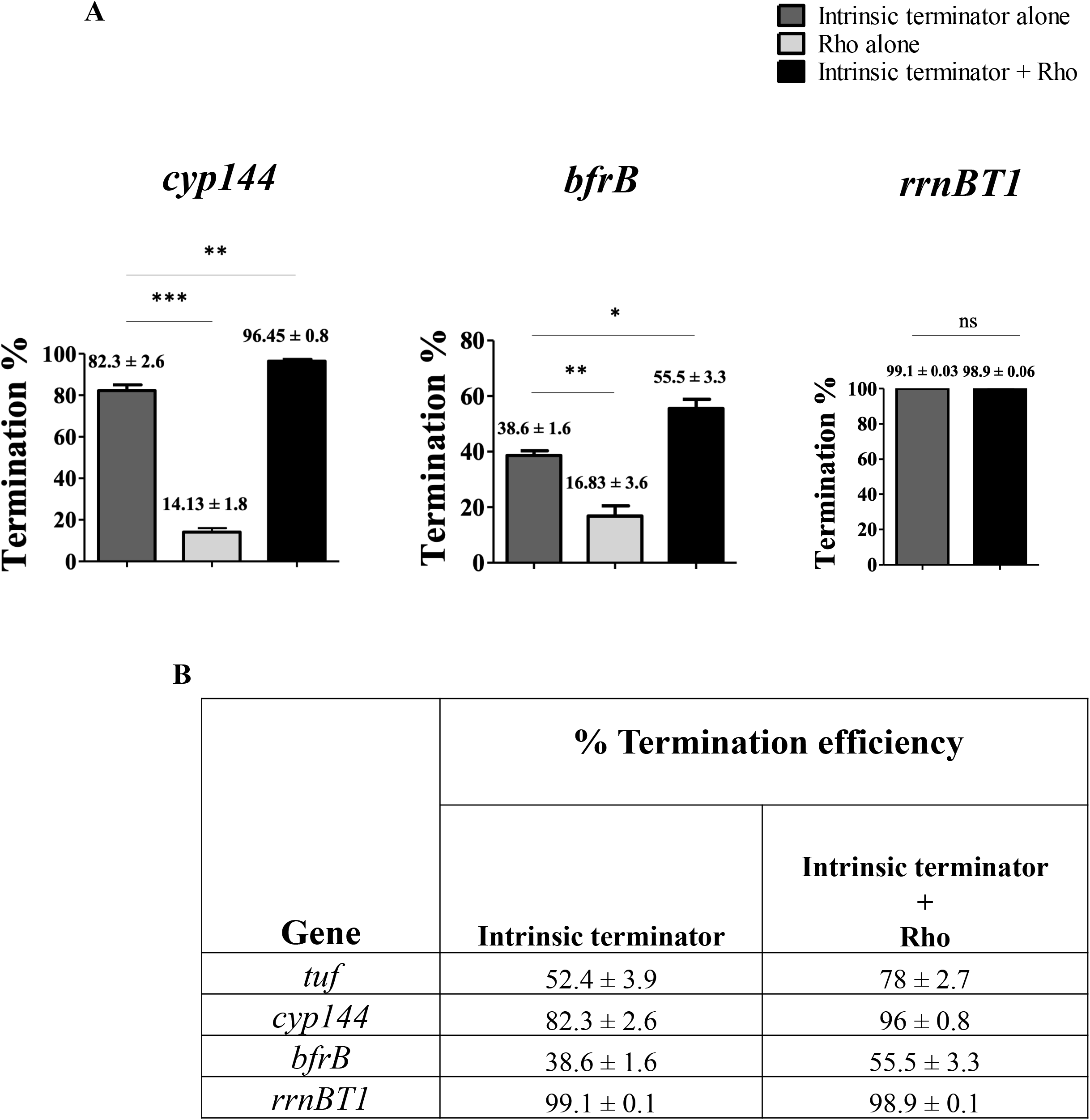
Transcription termination by intrinsic terminator and Rho at 3’UTR. **(A)** Assessment of transcription termination efficiency for *cyp144, bfrB*, and *rrnBT1* genes using the mRuby fluorescence readout as described in Material and methods. The data represents the contribution of intrinsic terminator and Rho individually and together in transcription termination. The mean ± SD is calculated from the three independent experiments. Statistics: One-way ANOVA followed by Tukey multiple-comparisons test, *p<0.05, **p<0.001,***p<0.0001, ‘ns’ is not significant. **(B)** The table depicts the impact of intrinsic terminator alone and intrinsic terminator + Rho together in transcription termination of *tuf, cyp144, bfrB*, and *rrnBT1*.

### Rho prevents transcriptional read-through in genes with weak intrinsic terminators

To verify the above results, transcription read-through was assessed by Real-Time-quantitative PCR (RT-qPCR) of the coding region and downstream UTR. The experimental strategy is depicted in Figure 3A and details are described in Materials and methods. We estimated the terminated and read-through transcripts of *bfrB* and *mkl* genes which have intrinsic terminators that lack U-tract (23). The levels of their transcripts were compared between wild type and Rho-CKD strain (Figure 3). By depleting Rho (Figure 3B and C), the quantity of transcripts increased at the 3’ UTR of *bfrB* and *mkl* genes (Figure 3D), pointing towards the role of Rho in suppressing transcriptional read-through at their 3’ ends. Thus, it appears that the two modes of transcription termination jointly contribute to prevent RNA polymerase (RNAP) continuing the unwanted RNA synthesis at the 3’ regions of these genes.

**Figure 3.**
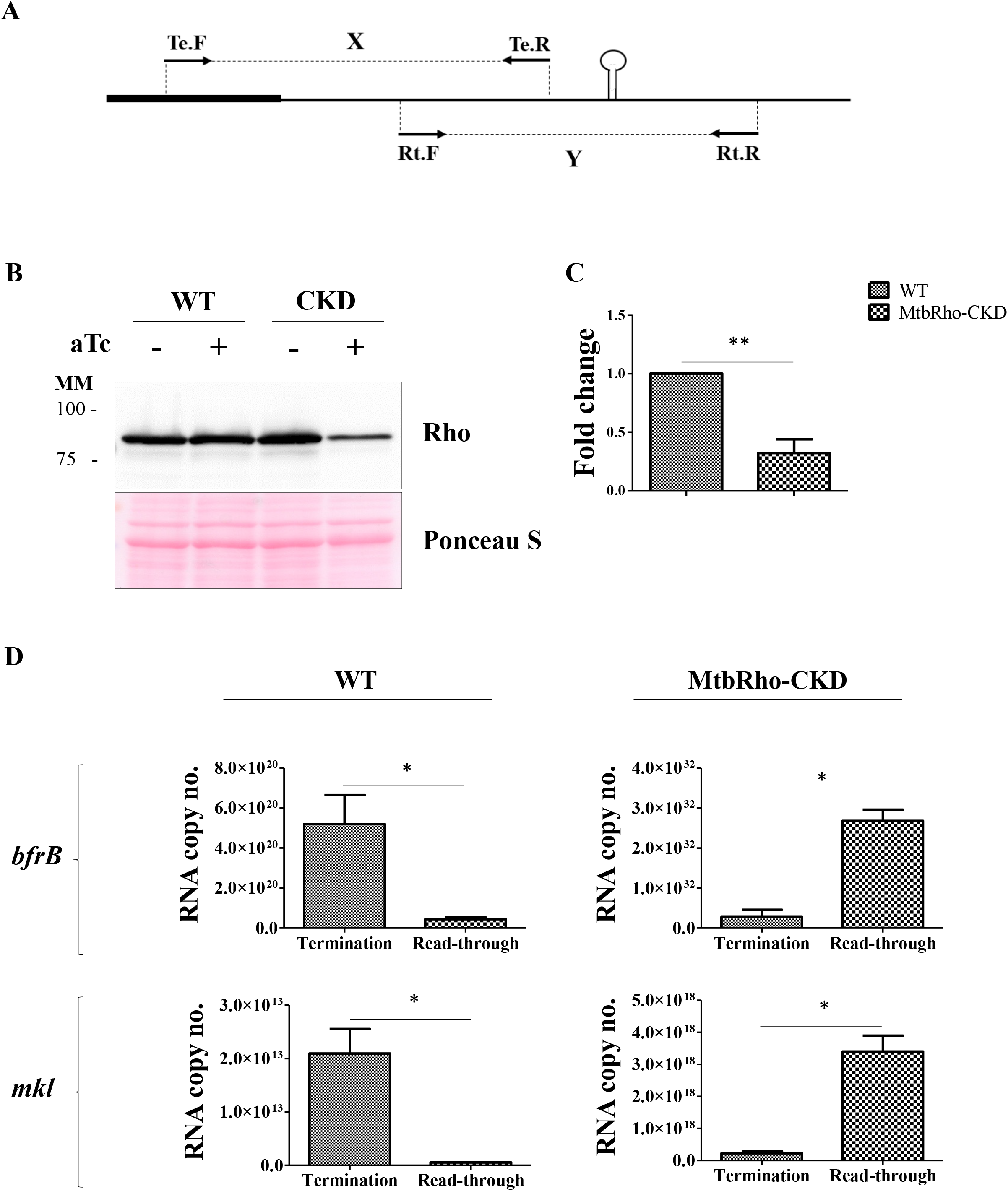
Rho suppresses transcriptional read-through at intrinsic terminators. **(A)** Schematic depiction of the RT-qPCR experiment to quantify the terminated (X) and read-through (Y) transcripts in genes carrying intrinsic terminator. Te.F-Te.R and Rt.F-Rt.R represent the primer sets to measure terminated and read-through transcripts respectively. **(B and C)** Comparison of protein and RNA level of Rho between wild type (WT) and Rho-CKD strain of MtbH37Rv by western blotting and RT-qPCR. MM represents molecular marker (kDa) and Ponceau S staining is shown as a loading control for western blotting. **(D)** Differential levels of the *bfrB* and *mkl* terminated and read-through transcripts between WT and Rho-CKD strains. The data represented is mean ± SD from the three independent experiments. The P-values were calculated by using unpaired, two-tailed t-test, *p<0.05, **p<0.005.

### Rho activity at the 3’ UTR of *mutT1, Rv3183, mkl, bfrB, metE, cyp144*, and *tuf*

Among the multiple steps involved in RDT, one of the crucial step is to initiate Rho’s action by its binding to the rut (rho utilization) site. *E. coli* Rho (EcRho) rut site essentially comprises C-rich, unstructured RNA (4, 33). Mycobacteria as such have G/C-rich genome and the nature of the rut site for MtbRho is not known. Our efforts to identify rut site have not provided any unambiguous sequence preference. Notably, the prediction tools employed to identify rut sites in *E. coli* and *Bacillus subtilis* (34, 35) do not provide satisfactory results for mycobacteria genomes. Due to their high G + C content there are far too many C-rich sites in their genomes. However, like *E. coli* and other Rho factors, MtbRho shows ATPase activity with polycytidylic acid (poly C) substrates (Supplementary Figure 4). To examine whether MtbRho activity can be detected in the RNA substrates used above, ATPase assays were carried out with the enzyme using *in vitro* transcribed 3’ UTR of *mutT1, Rv3183, mkl, bfrB, metE, cyp144*, and *tuf* genes. The results presented in Figure 4 show that MtbRho can hydrolyse ATP in the presence of these RNA substrates. The ATPase activity was specific to Rho as it was inhibited by bicyclomycin (BCM). A deletant, ΔWQRRho which lacks Walker B motif, Q-loop, and R-loop of the ATPase domain showed no ATP hydrolysis (Figure 4). These results suggest that MtbRho can use RNA substrates harboring an intrinsic terminator. It also implies that, the intrinsic terminator in the transcript does not negatively impact RDT.

**Figure 4.**
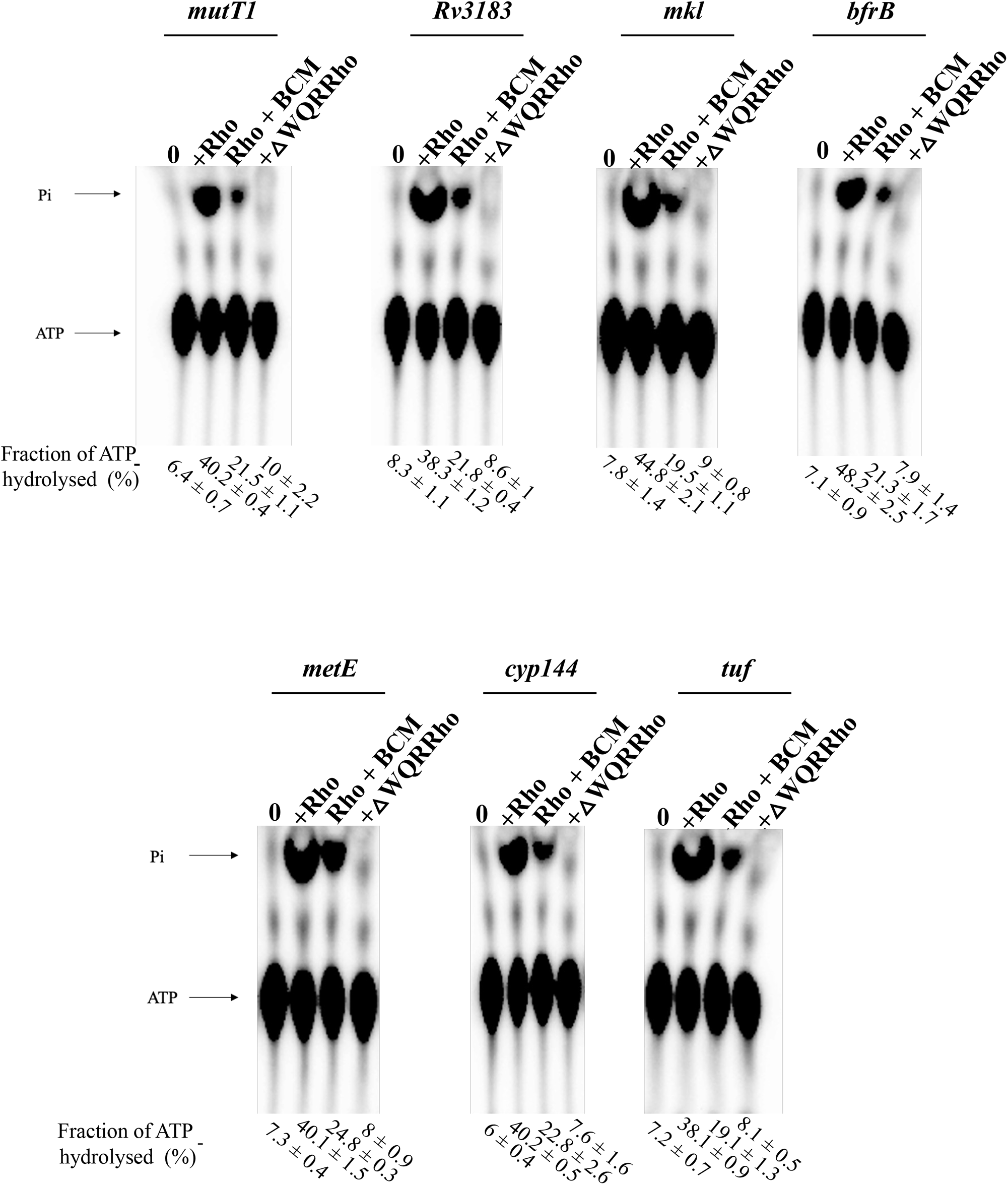
ATP hydrolysis by MtbRho with RNA carrying intrinsic terminator. MtbRho (100 nM) hydrolyses ATP in presence of *in vitro* transcribed RNA (300 µM) containing intrinsic terminators. ATP hydrolysis is inhibited in the presence of 600 µM bicyclomycin (BCM). ATPase activity was not seen with ΔWQRRho (100 nM). The released inorganic phosphate (Pi) was visualized using Phosphorimager. Percentage fraction of Pi release was calculated and SDs were determined from three independent experiments.

### Rho-dependent transcription termination augments intrinsic termination

From the results in Figure 1-3, it is apparent that Rho imparts its effect downstream of the 3’ end of genes that have intrinsic terminator with suboptimal or no Us, and transcription termination *in vivo* could be a culmination of contribution by Rho. To investigate the participation of Rho in transcription termination in templates containing intrinsic terminators, *in vitro* transcriptions were carried out with Msm RNAP and templates having T7A1 promoter (Figure 5A). The promoter is often employed for *in vitro* transcription reactions as it is, i) well-characterized (36–39); ii) already used for studying transcription termination(34, 40, 41); and iii) recognized and transcribed efficiently by both Msm and Mtb RNAPs, besides *E. coli* RNAP (42, 43). In the *in vitro* reactions, termination efficiency solely depends on the strength of an intrinsic terminator in absence of Rho. When Rho is present, the additional contribution by the factor can be assessed.

**Figure 5.**
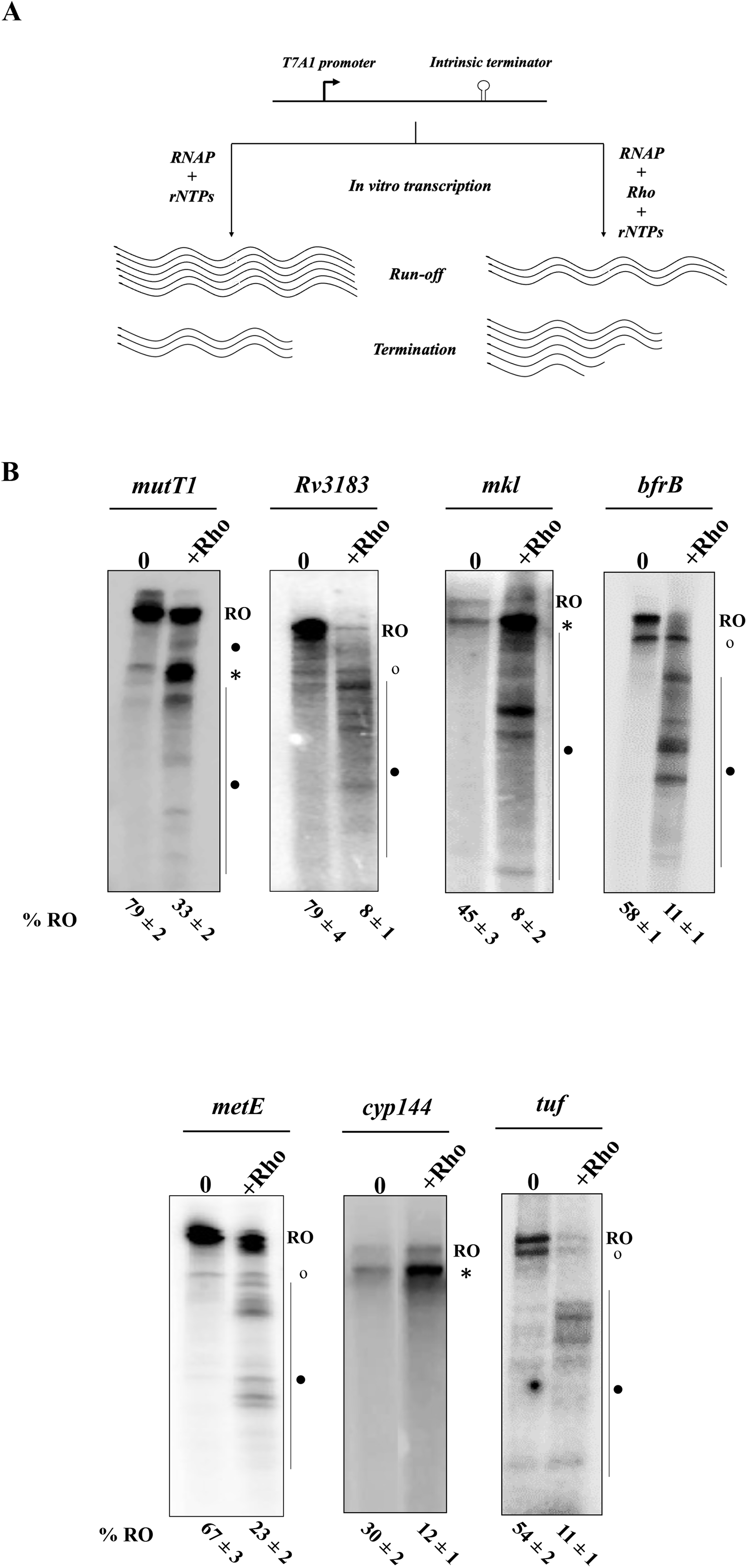
Intrinsic terminator and Rho interact for transcription termination. **(A)** Schematic representation of *in vitro* transcription strategy to study intrinsic terminator and Rho functional interaction. **(B)** *In vitro* transcription assays were carried out using 10 nM linear DNA templates harboring intrinsic terminator (*mutT1, Rv3183, mkl, bfrB, metE, cyp144*, and *tuf*) downstream of T7A1 promoter in the presence and absence of MtbRho (100 nM) as described in Materials and methods. Transcripts were resolved on 6% 8M urea-PAGE. RO corresponds to run-off transcripts. Intrinsic termination products are marked with open circles (∘). Rho-dependent termination products are marked with closed circle (•). Asterisk (*) represents the signal corresponding to the termination event by both intrinsic terminator and Rho together. Run-off percentage (% RO) were calculated (using the formula : (Run-off)/(Run-off +Total termination) and represented with SD from three experimental replicates (shown below the gel lanes).

In these assays, MtbRho substantially increased the transcription termination for all the tested templates (Figure 5B). In templates containing *cyp144* intrinsic terminator, RDT was observed at the site of intrinsic termination. With *mutT1, and mkl* templates, RDT was observed downstream of the stop codon at different locations in addition to the intrinsic termination site. With *mkl*, RDT signals were found between the stop codon and the intrinsic terminators. However, with *mutT1*, RDT signals were also seen on either side of the intrinsic terminator, indicating wide landscape in the 3’ UTR for Rho action. In the case of *Rv3183, bfrB, metE*, and *tuf* templates, intrinsic termination and RDT signals appeared at independent locations, with RDT preceding the structure. Nevertheless, their additive impact resulted in increased termination of transcription at these templates (Figure 5B). In none of these templates RDT signals were found exclusively downstream of intrinsic terminators, although such pattern has been described in *E. coli* (16, 32). Rho inhibitor BCM inhibits MtbRho activity albeit at high concentrations (21). BCM reduced the RDT signals without affecting the intrinsic termination in these reactions (Supplementary Figure 5A). In assays with Rho mutant ΔWQRRho, no change in termination efficiency was observed; termination pattern resembled those reactions where Rho was not added (Supplementary Figure 5A). To verify whether these RDT events result in the dissociation of transcripts, transcript release experiments were carried as described (23, 24). In assays with streptavidin bead-bound DNA templates (see Materials and methods), transcript release was observed in the supernatant fraction. The fraction of terminated transcripts increased in the presence of MtbRho (Supplementary Figure 5B).

### Intrinsic terminator and Rho collaborate for efficient transcription termination in *E. coli*

Generally, the two modes of transcription termination are treated as stand-alone processes in *E. coli*. To assess the participation of both intrinsic terminator and Rho together, the above-described assays were performed in *E. coli*. mRuby fluorescence and *in vitro* transcription assays were carried out with *E. coli* intrinsic terminators in the presence and absence of Rho. In addition to *rrnBT1*, previously identified *E. coli* intrinsic terminators *aroG* and *ycbL* (16) were chosen for these assays. These intrinsic terminators have a relatively weaker stem (ΔG = −13 kcal/mol) compared to the strong terminators (ΔG ≤ −25 kcal/mol) and they are followed by 8 U’s (12, 16, 44). The strength of these intrinsic terminators was assessed *in vivo* as the percentage of mRuby fluorescence in EcRho temperature-sensitive strain AMO14 (45) and knockout (KO) strain RS1309 (46). The deletion of rac (Δrac) ensures Rho-KO survival in RS1309 in which wild-type Rho can be expressed from an inducible plasmid (46). In these strains, mRuby expression was higher in the absence of Rho and/or when intrinsic terminator was not introduced (Figure 6A and Supplementary Figure 6A). However, when *aroG* and *ycbL* intrinsic terminators were introduced, mRuby expression was markedly reduced. Further reduction in the fluorescence signal in the presence of Rho (At permissive temperature for AMO14 strain and IPTG induction of Rho introduced through a plasmid to RS1309; see Materials and methods) suggest that transcription at the 3’ end of *aroG* and *ycbL* genes is decreased by the additive effect of their intrinsic terminator and Rho (Figure 6B and Supplementary Figure 6B). In contrast, *rrnBT1* terminator no dependence on Rho for termination; termination efficiency was unaltered in the presence and absence of Rho (Figure 6B and Supplementary Figure 6B). The difference in the mRuby fluorescence level in these two *E. coli* strains before the introduction of terminators appears to be due to the strain difference. While mRuby expression is driven by hsp60 promoter in both the strains, Rho induction in RS1309 is by IPTG, whereas in AMO14 it is thermo inducible. As a result high mRuby expression is seen in AMO14.

**Figure 6.**
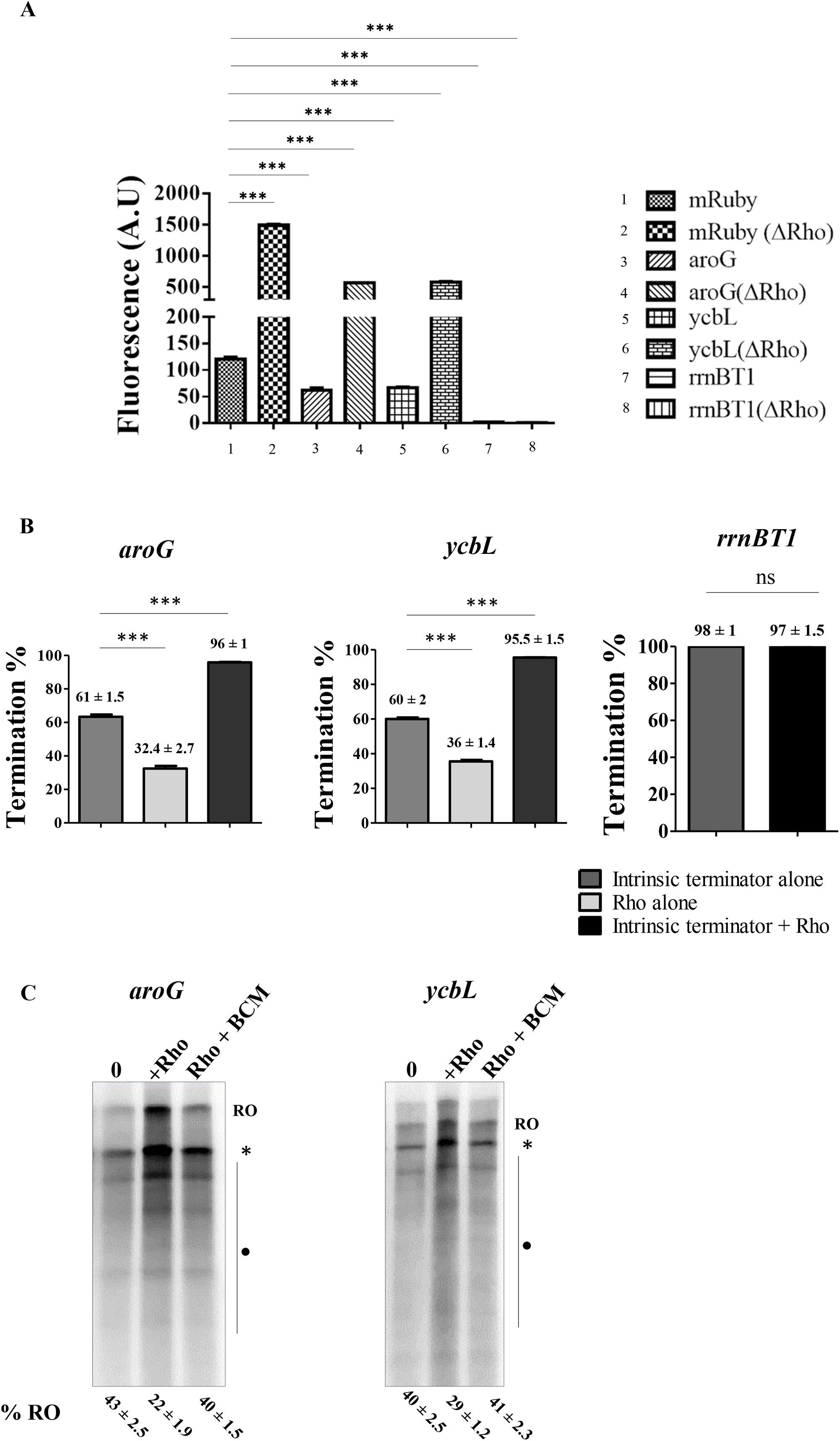
Measurement of contribution of intrinsic terminator and Rho in *E. coli*. **(A)** Comparison of mRuby expression level for *aroG, ycbL*, and *rrnBT1* intrinsic terminators, in the presence and absence of Rho, in *E. coli* AMO14 strain. **(B)** Calculation of termination efficiency and contribution of intrinsic terminator-alone, Rho-alone and intrinsic terminator + Rho in transcription termination of *aroG, ycbL*, and *rrnBT1* genes using mRuby fluorescence readout. The data represented is mean ± SD from the three independent experiments. The P-values were calculated by one-way ANOVA followed by Tukey multiple-comparisons test,***p<0.0001, ns – not significant. **(C)** *In vitro* transcription assays were carried out using 10nM linear DNA templates harboring *aroG* or *YcbL* intrinsic terminator, *E. coli* RNAP (200 nM) in the presence and absence of EcRho (100 nM) and BCM (300 µM) as described in Materials and methods. Transcripts were resolved on 6% 8M urea-PAGE. Rho-dependent termination products are marked with closed circle (•). Asterisk (*) represents the signal that corresponds to the termination event by both intrinsic terminator and Rho together. RO marks the run-off transcripts. Run-off percentage (%RO) were calculated and represented with SD from three experimental replicates (shown below the gel lanes).

*In vitro* transcription assays carried out with *E. coli* RNAP and linear templates carrying *aroG* and *ycbL* intrinsic terminators transcribed by the T7A1 promoter showed a substantial increase in the termination at the site of intrinsic termination when EcRho was added (Figure 6C). The presence of BCM reduced the RDT signals. Together, the results from the *in vivo* fluorescence assays and *in vitro* transcription assays indicate that intrinsic terminator and Rho collaborate to prevent unwanted transcriptional read-through in *E. coli* as well.

## Discussion

We describe participation of both the modes of transcription termination wherein intrinsic terminators and Rho function together for a more effective stop of RNA synthesis in a given 3’ UTR. The need for such collaboration is apparent as a majority of intrinsic terminators are inefficient in terminating transcription (22–25). The interplay of the two processes seems necessary to enhance the overall efficiency of termination at the 3’ end of the genes. Such an arrangement would ensure the control of undesired transcriptional read-through into downstream genes. Transcriptional read-through can affect gene expression by increasing transcription of downstream genes oriented in the same direction, destabilizing transcripts from convergent transcription units, and/or changing the expression of global regulators causing genomic instability (10, 25, 31, 47). The *in vitro* transcription and *in vivo* reporter assays described here show that Rho participates in transcription termination at the intrinsic terminators or in the neighbouring regions. Inadequate levels of Rho in Rho-CKD cells allowed the transcription read-through, highlighting the necessity of Rho in impacting the process.

Although individually the two mechanisms of transcription termination are well illustrated, their combined action, and *in vivo* interplay was not given much consideration. Studies that mapped the 3’ ends of RNA revealed the role of the nucleases in masking the actual Rho-mediated termination sites at the 3’UTRs (14, 37). While these end-mapping in Rho or nuclease background indicated the contribution in the termination processes, the mRuby reporter system described here directly examines the extent of participation of the two processes individually and together. High stability and resistance to denaturation at extreme conditions makes mRuby a suitable choice for these *in vivo* assays for quantitative assessment of individual and collective contribution by the two mechanisms (48–50). Although the system is used for mycobacteria and *E. coli* here, the ease of its application to other bacteria should facilitate further studies.

The textbook definition of intrinsic terminators having a strong G/C-rich hairpin followed by 7-9 U residues (29) are found only in the strongest of terminators such as rRNA (22) and hence they function efficiently on their own both *in vitro* and *in vivo*. Now it is apparent that suboptimal and weak terminators are found in *Bacillus* (25, 51), *E. coli* (37), mycobacteria (23, 24) and other species (14, 26). In *E. coli*, a significant fraction of the hairpin loop structures present at the 3’ end of the genes lack a discernible U trail (14). Moreover, in various bacteria, the U trail has been shown to be either essential (52, 53), unnecessary (54), or necessary in specific conditions (55). Variation in the length of the stem, its strength (ΔG), and the number of U residues of any intrinsic terminator can influence its termination efficiency (10, 50, 52, 54). The physiological conditions in a bacterium (including RNAP elongation rate and NTP availability) are also crucial factors in determining the termination efficiency (57, 58). Hence, we extended our studies to *E. coli* to see whether intrinsic terminator and Rho functional interaction is a more widespread phenomenon. *In vitro* and *in vivo* experiments with *E. coli* intrinsic terminators and Rho also revealed their cumulative effect to downregulate transcriptional read-through (Figure 6 and Supplementary Figure 6).

With their distinct mode of termination, one requiring a specific secondary structure and the other relying on specific protein-RNA interactions, the two conserved mechanisms have been generally regarded as stand-alone mechanisms (1, 4, 33, 59, 60). However, terminator inefficiency and the consequent read through (22–25) (Supplementary Table S2) pointed at the contribution of additional factors *in vivo*. Increase in termination efficiency by NusA in *in vitro* reactions has been observed in *E. coli* (61, 62) and the mechanism elucidated (63). Now, the role of NusA and/or NusG have been examined in *B. subtilis*, Mtb, and *Mycobacterium bovis* (24, 25, 51, 64). Suboptimal intrinsic terminators having lower number of U residues downstream of the stem serve as substrates for NusA and NusG in *B. subtilis* (25, 51). Notably, a subset of these terminators having non-canonical architecture are dependent on NusA (25) to bring about efficient termination both *in vitro* and *in vivo*. In the terminators where NusG is involved, its role is in pausing of RNAP whereas NusA is suggested to have a role in hairpin folding and/or stabilization (23, 46). In contrast to the role of NusG in *B. subtilis*, in Mtb, NusG does not seem to enhance termination (64), while in *M. bovis*, it facilitates termination at non-canonical terminators (24). As NusA and NusG are already known to modulate intrinsic termination (3, 25, 51, 65, 66), contribution from other factors in improving the termination efficiency is not unanticipated. Indeed, in *B. subtilis* Rho null mutant, transcription read-through from inefficient intrinsic terminators of antisense transcripts was observed (67). Rho participation described here is likely to be at a step after the formation of hairpin and pause based on prevailing understanding of Rho action (4, 33, 40, 60, 68). Apart from the bonafide RDT sites, intrinsic terminator induced pausing of RNAP with or without the contribution of Nus factors can also facilitate Rho to act and terminate transcription at the intrinsic termination sites.

Given their suboptimal architecture, many stem-loop structures were thought not to function as terminators, and act as pause sites during transcription (69). An additional role attributed to them is in protecting the 3’ end of mRNA from the action of polynucleotide phosphorylase (PNPase), RNaseII and possibly other nucleases (16, 32). In one of the RNA 3’ ends mapping studies the structures were shown to protect from exonuclease action subsequent to RDT downstream (16). While they suggested that the 3’ protected ends could serve as intrinsic terminators, the possibility was not examined (16, 32). In the other study, it was shown that *gal* mRNA is subjected to both intrinsic termination and RDT. The suboptimal intrinsic terminator of *gal* operon doubled up to protect the 3’ end from exonucleases, masking the detection of RDT downstream (32). Together the two mechanisms contribute to more effective termination at *gal* operon (32). Although it was suggested that the two modes act independent of each other, from our data it is apparent that the signals are likely to be connected for better control of gene expression in some of the 3’ UTRs.

Based on the present study, we considered different patterns of Rho action beyond the stop codons of the genes having weak intrinsic terminators (Figure 7) 1) intrinsic termination and RDT at distinct locations with RDT occurring first; 2) both at the same site; 3) RDT at the site of intrinsic termination and before; 4) RDT before, after, and at the intrinsic termination site; 5) RDT both at the site of intrinsic termination and downstream; 6) intrinsic termination and RDT at distinct locations with intrinsic termination occurring first. Among these possibilities, the data presented (Figure 5) provide examples for the first four with mycobacterial intrinsic terminators and MtbRho. The 6^th^ pattern was described by Wang et al (32). The 5^th^ pattern is likely to be found if more terminators are analysed given that in many phyla across the bacterial domain, the intrinsic terminators do not always confine to the first 30 nucleotides after the stop codon; the stop-to-stem distance can range from 0-150 nt (12, 20, 26, 58). Notably, the prevailing two modes of Rho action can account for its participation in the patterns described (4, 40, 60, 68). Rho could translocate to catch up with the paused RNAP at the weak intrinsic terminator and induce conformational change in the enzyme to terminate transcription. Alternately, pre-formed Rho-RNAP complex can form a pre-termination complex at the terminator to allosterically inactivate the RNAP.

**Figure 7.**
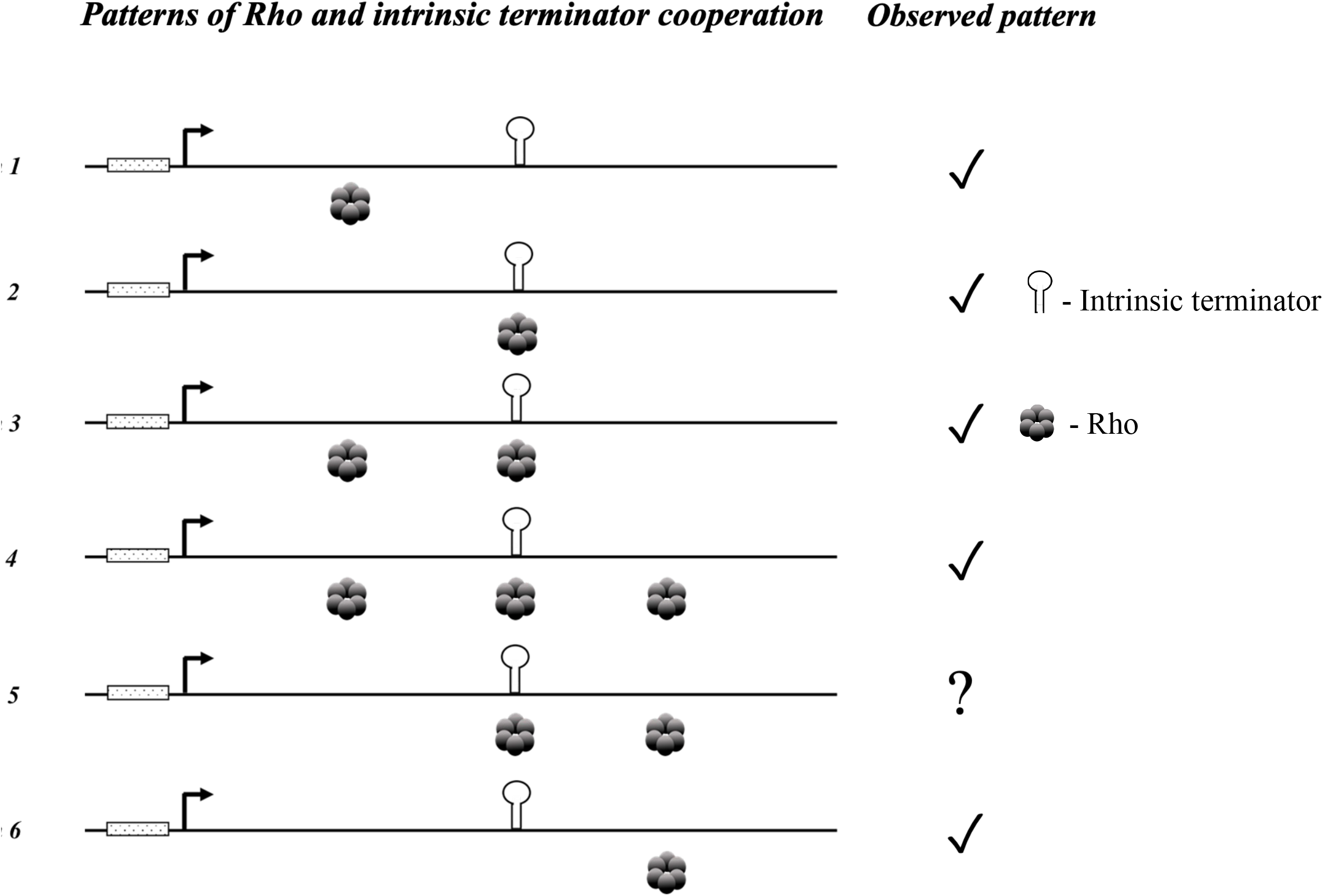
Different modes of Rho and intrinsic terminator functional interaction. Schematic depiction of various Rho and intrinsic terminator interactions in transcription termination. Rho and intrinsic terminator positions in the figure depicted represent their site of action at the UTR. Tick mark (✓) highlights the Rho – intrinsic terminator cooperation pattern observed so far. Question mark indicates lack of examples for the pattern.

To conclude, the study of transcription termination has come a full circle. From the emerging theme it is apparent that even in the well-studied systems such as *E. coli* and *B. subtilis* intrinsic termination can be achieved with non-canonical structures. Although initially described in mycobacteria, it appears that non-canonical intrinsic terminators are found across bacterial kingdom. In many of these sites, a more efficient termination is brought about by the collaboration with Rho or other elongation factors like NusA and NusG. It remains to be seen whether Rho can combine with other elongation factors to ensure transcriptional read-through is curtailed. The association of NusA, NusG, and Rho with RNAP in recently elucidated structures (40, 60), should facilitate in understanding their combined function in transcription elongation and termination.

## Materials and methods

### Bacterial strains, culture conditions, plasmids, chemicals and enzyme

Mtb H37Rv, Msm mc^2^155, *E. coli* AMO14 (ts) (45) and RS1309 (Δrho Δrac) (46) strains were used for transcription termination studies. *E. coli* DH10B and BL21 λDE3 were used for cloning and protein overexpression, respectively (Supplementary Table S3). Msm and Mtb were cultured in Middlebrook 7H9 broth supplemented with 0.5% albumin, 0.2% dextrose, 0.085% NaCl, 0.2% glycerol and 0.05% Tween 80, and with antibiotics when appropriate (50 μg/ml hygromycin and 25 μg/ml kanamycin). Mtb and Msm Rho-CKD strains were grown at 37°C and 30°C respectively with shaking at 120 rpm. For induction of small guide RNA (sgRNA) and *dcas9* expression from pRH2521 and pRH2502 plasmids respectively (70), Msm and Mtb Rho-CKD cultures were supplemented with anhydrotetracycline (aTc) to a final concentration of 200 ng/ml. The addition of aTc was repeated every 48 h to maintain induction of *dcas9* and sgRNAs when required. When grown on Middlebrook 7H11 agar supplemented with 0.5% glycerol and 10% oleic acid-albumin-dextrose-catalase, aTc (100 ng) was added on a paper disc when indicated. *E. coli* AMO14 and RS1309 strains were grown in LB broth having appropriate antibiotics (45, 46). Kanamycin (50 μg/ml) and chloramphenicol (30 μg/ml) were used for AMO14 (45). Tetracycline (30 μg/ml), ampicillin (100 μg/ml), and IPTG (1mM) were added for EcRho expression in RS1309 (46).

DNA fragments carrying intrinsic terminators were cloned in pUC18 plasmid downstream of the T7A1 promoter (Supplementary Table S1) (23). Oligonucleotides were synthesized by Sigma (Supplementary Table S4). α^32^P UTP was purchased from the Board of Radiation and Isotope Technology (BRIT), Mumbai. DNA modifying enzymes were purchased either from NEB or Roche. Streptavidin-coated agarose beads were from GE healthcare. Sigma A (SigA) enriched His-tagged RNAP was purified from Msm SM07 strain as described before (71) and was used for *in vitro* transcription assays. The enzyme (Msm RNAP) was purified by polyethyleneimine precipitation, heparin sepharose chromatography, and Ni-NTA affinity chromatography (71). MtbRho and its mutant that lacks the Walker B motif, Q-loop, and R-loop (ΔWQRRho) of the ATPase domain were purified by polyethyleneimine and ammonium sulphate precipitation, elution from Heparin and SP-sepharose chromatography columns as described previously (72). EcRho with a C-terminal His-tag was expressed in *E. coli* BL21 λDE3 cells. The lysate was passed through the Ni-NTA column, eluted with 500 mM imidazole, and further purified using the Hitrap-heparin column (72).

### Construction of Rho-CKD strains

pRH2502, a vector expressing an inactive version of *Streptococcus pyogenes cas9 (dcas9)*, and pRH2521, a vector to express the gene-specific sgRNA were obtained from Robert N. Husson (70). To generate Rho-CKD in Msm and Mtb, Rho-specific sgRNAs were designed (Supplementary Figure 1A) and cloned in pRH2521 vector. Protospacer adjacent motif (PAM) sequence was identified in the proximal coding region for both the strains of mycobacteria. These vectors were transformed into Msm/Mtb cells containing pRH2502 (for *dcas9* expression).

### mRuby fluorescence assay to estimate transcription termination *in vivo*

DNA fragment containing the *mRuby* gene (48) was cloned in the pMV261 vector (73) downstream hsp60 promoter between EcoRI and HindIII sites. Intrinsic terminators from *tuf, cyp144* and *bfrB* genes of Mtb were cloned upstream of *mRuby* between BamHI/PvuII and EcoRI sites (Supplementary Table S1 and S3). The DNA fragment containing hsp60 promoter-intrinsic terminator-*mRuby* was excised from pMV261 and cloned in pRH2521 vector at NotI and HpaI sites (Supplementary Table S1). This pRH2521 vector contains MsmRho specific sgRNA. Msm cells carrying pRH2502 vector expressing dCas9 protein were electroporated with pRH2521-*mRuby*/intrinsic terminator*-mRuby* and colonies from each of these strains were grown till O.D 0.5 in Middlebrook-7H9 media at 30°C (Supplementary Figure 2A). Cultures were washed and resuspended in 1X phosphate-buffered saline. The fluorescence readouts were normalized against the O.D. 300μl of 0.5 O.D culture from these strains were used to measure mRuby fluorescence. The fluorescence was estimated using excitation and emission wavelength of 558nM and 605nM, respectively in the infiniteM200PRO-TECAN.

DNA fragments containing intrinsic terminators *aroG, ycbL* and *rrnBT1* from *E. coli* K-12 MG1655 were cloned into pRH2521 vector upstream of *mRuby* gene and transformed into *E. coli* AMO14 and RS1309 strains. mRuby fluorescence assays were carried out by growing respective test and control strains in LB media until 0.5 O.D. at 37 °C. For EcRho-KO strain RS1309, mRuby fluorescence was measured at the culture density of 0.5 O.D. For the temperature-sensitive *E. coli* AMO14 strain, where the genomic *Rho* gene is inactivated and a functional copy of *E. coli Rho* is supplied on a temperature-sensitive plasmid (45), mRuby fluorescence assay was performed after shifting and incubating the cultures to non-permissive temperature (42 °C) as described (45).

### Western blot analysis

The concentration of proteins from the lysates of the exponential phase cultures of Mtb and Msm were estimated by the Bradford method for both the wild type and Rho-CKD strains, in the presence and absence of aTc. Proteins were separated on 8% SDS-PAGE. Rho was detected by immunoblotting with 1/20000 dilutions of anti-MtbRho antisera and visualized by chemiluminescence. Blots were also stained with Ponceau S to depict the protein in each lane.

### RNA isolation and RT-qPCR

RNA was isolated by using RNAZol-RT reagent (Sigma) from exponentially growing MtbH37Rv/Msm mc^2^155 and its Rho CKD strain. cDNA was synthesized by the Applied Bioscience cDNA synthesis kit following the manufacturer’s instructions using the gene-specific reverse primer for *bfrB* and *mkl* and random primers for MtbRho, MsmRho, mRuby, and Hyg^R^. RT-qPCR was carried out in a BioRad CFX96 Touch Real-Time PCR Machine. To quantify the terminated and read-through transcripts, two primer pairs were designed targeting upstream and downstream region of the *bfrB* and *mkl* intrinsic terminators (23, 74). A standard curve was generated using the known concentration of *bfrB* and *mkl* RNA that was *in vitro* transcribed using T7 RNAP in the presence of 2 mM rNTPs (Supplementary Figure 7 and Supplementary Table S1). Terminated and read-through transcripts of *bfrB* and *mkl* control and MtbRho-CKD strains were interpolated from the standard curve (Figure 3D). RT-qPCR to compare the RNA levels of MtbRho, MsmRho, mRuby, and Hyg^R^ were carried out for control and Rho-CKD strains. Quantitation of Hyg^R^ in various conditions served as an indicator of plasmid copy number (75, 76). 16SrRNA was used as an internal control to estimate the fold change (2^-ΔCT^) in the levels of MtbRho, MsmRho, mRuby, and Hyg^R^ transcripts in the above strains.

### ATPase assays

ATPase assays were carried with poly C (40 ng) or *in vitro* transcribed 3’UTR of *mutT1, Rv3183, mkl, bfrB, metE, cyp144*, and *tuf* (300 µM) by T7 RNAP in the presence of 2 mM rNTP (Supplementary Table S1). RNA substrates were incubated in T-Buffer (Tris HCl pH8 50 mM, MgOAc 3 mM, potassium glutamate 100 mM, DTT 0.1 mM, EDTA 0.1 mM, BSA 0.1 mg/ml, Glycerol 5%) in the presence of 100 nM MtbRho with or without 600 µM BCM or ΔWQRRho. The reactions were initiated by the addition of 1 mM of ATP followed by incubation at 37°C for 30 min. 100 nCi of γ^32^P-ATP was used as a tracer in the radiometric assay to detect the formation of inorganic phosphate (Pi); the reactions were stopped by adding chloroform. Following centrifugation, the aqueous phase was used to estimate the release of Pi by resolving on polyethyleneimine thin-layer chromatography sheets using LiCl (1.2 M) and EDTA (0.1 mM) as the mobile phase and visualized with the Typhoon 9500(GE) Phosphorimager.

### *In vitro* transcription

Intrinsic terminators of *mutT1, Rv3183, mkl, bfrB, metE, cyp144*, and *tuf* genes of Mtb, and from *E. coli*, intrinsic terminators of *aroG* and *ycbL* genes were selected for *in vitro* transcription assays (Supplementary Table S1). DNA templates for transcription reactions were prepared as described (23). 10 nM DNA template and 200 nM RNAP from Msm or *E. coli* were incubated in T-Buffer, first on ice followed by 10 min at 37°C. MtbRho or EcRho (100 nM) were added to the reaction mixture and incubated at 37°C for 5 min. When BCM (600 µM) was used, it was pre-incubated with Rho before adding to the reaction mixture. Transcription was initiated with the addition of 200 µM rNTPs (200 µM mix of rATP, rCTP, rGTP, and 20 µM rUTP) and 10µCi α-^32^P-UTP, incubated at 37°C for 30 min and terminated by the addition of phenol-chloroform. Following centrifugation, an equal volume of gel-loading dye (95% deionized Formamide, 0.05% Bromophenol Blue, and 0.05% Xylene cyanol) was added to the aqueous phase, heated at 90°C for 2 min, and resolved on 8% 8M urea-PAGE. The gels were scanned and quantified using Typhoon 9500(GE) Phosphorimager and Multi Gauge V2.3 software respectively. The extent of combined transcription termination by intrinsic terminator and Rho was assessed by comparing the run-off levels in the presence and absence of Rho.

### Termination assays using immobilized DNA templates

Assays were carried out using immobilized DNA templates as described (23). Reactions were set up in duplicates, templates were generated by PCR amplification using biotinylated pUC18 forward primer. 10 nM biotinylated templates were incubated with streptavidin-coated agarose beads preequilibrated with T-buffer as described at 37°C for 15 min. *In vitro* transcriptions were carried out in the presence and absence of 100 nM MtbRho at 37°C for 30 min. One set was separated into supernatant and beads before processing and the other set was directly processed as described (23).

## Data Availability

The authors confirm that all relevant data has been provided either in the article or as Supplementary Material.

## CRediT Author Contribution

**Ezaz Ahmad:** Conceptualization, Methodology, Investigation, Visualization, Writing-Original draft, **Varsha Mahapatra:** Resources and Investigation, **Vanishree V M:** Resources and Investigation, **Valakunja Nagaraja:** Conceptualization, Supervision, Writing - Review & Editing, Funding acquisition.

## Acknowledgements

We thank Robert N. Husson, for pRH2521 and pRH2502 vectors, J. P. Richardson for *E*.*coli* AMO14 and Ranjan Sen for *E*.*coli* RS1309. We thank Umesh Varshney for careful reading of the manuscript and for providing mRuby vector. We acknowledge Rajiv Kumar Jha and Anirban Mitra for *E. coli* RNAP and EcRho respectively and the central facility of phosphor imaging of Indian Institute of Science, supported by Department of Biotechnology (DBT), Government of India.

## Abbreviations

RDT: Rho-dependent termination
TEC: transcription elongation complex
UTR: Untranslated region
CKD: Conditional knockdown
KO: Knockout
Mtb: *Mycobacterium tuberculosis*
Msm: *Mycobacterium smegmatis*
RT-qPCR: Real-Time-quantitative PCR
RNAP: RNA dpolymerase
rut: rho utilization
EcRho: *E. coli* Rho
BCM: Bicyclomycin
PNPase: Polynucleotide phosphorylase
sgRNA: Small guide RNA
PAM: Protospacer adjacent motif
aTc: Anhydrotetracycline
Hyg^R^: Hygromycin resistant gene
Pi: Inorganic phosphate;

