## Supplementary figures for "Intrinsic and Rho-dependent termination cooperate for efficient transcription termination at 3’ untranslated regions"

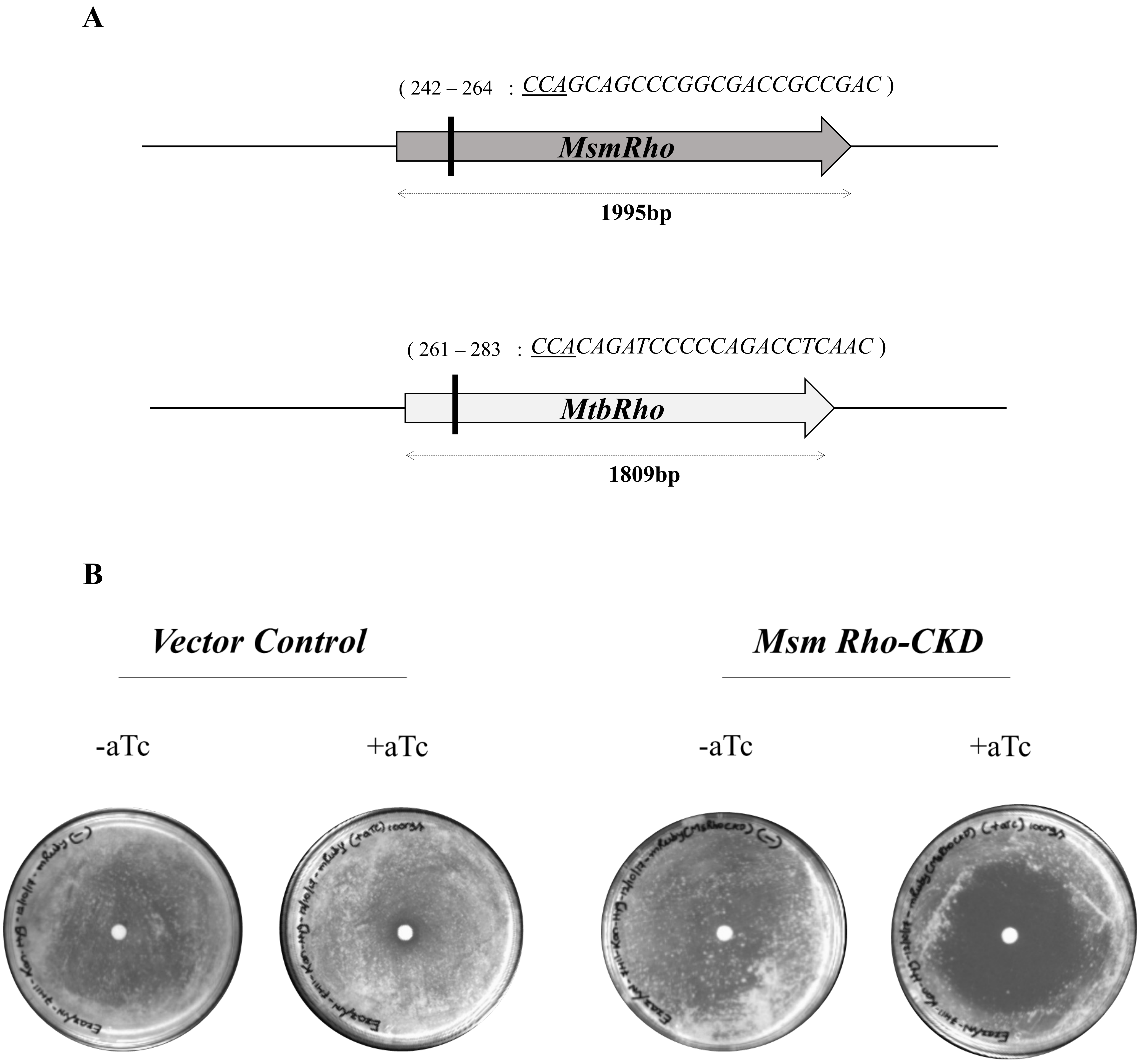

**Supplementary Figure 1. Rho depletion inhibits growth in mycobacteria (A)** Schematic depicting position and sequence of proximal guide RNA designed to generate CRISPRi mediated Rho-CKD in Msm and Mtb. **(B)**. Depletion of Rho level by aTc induction inhibits the growth of Msm, 100 ng of aTc was added on a disc placed at the center of the agar plate.

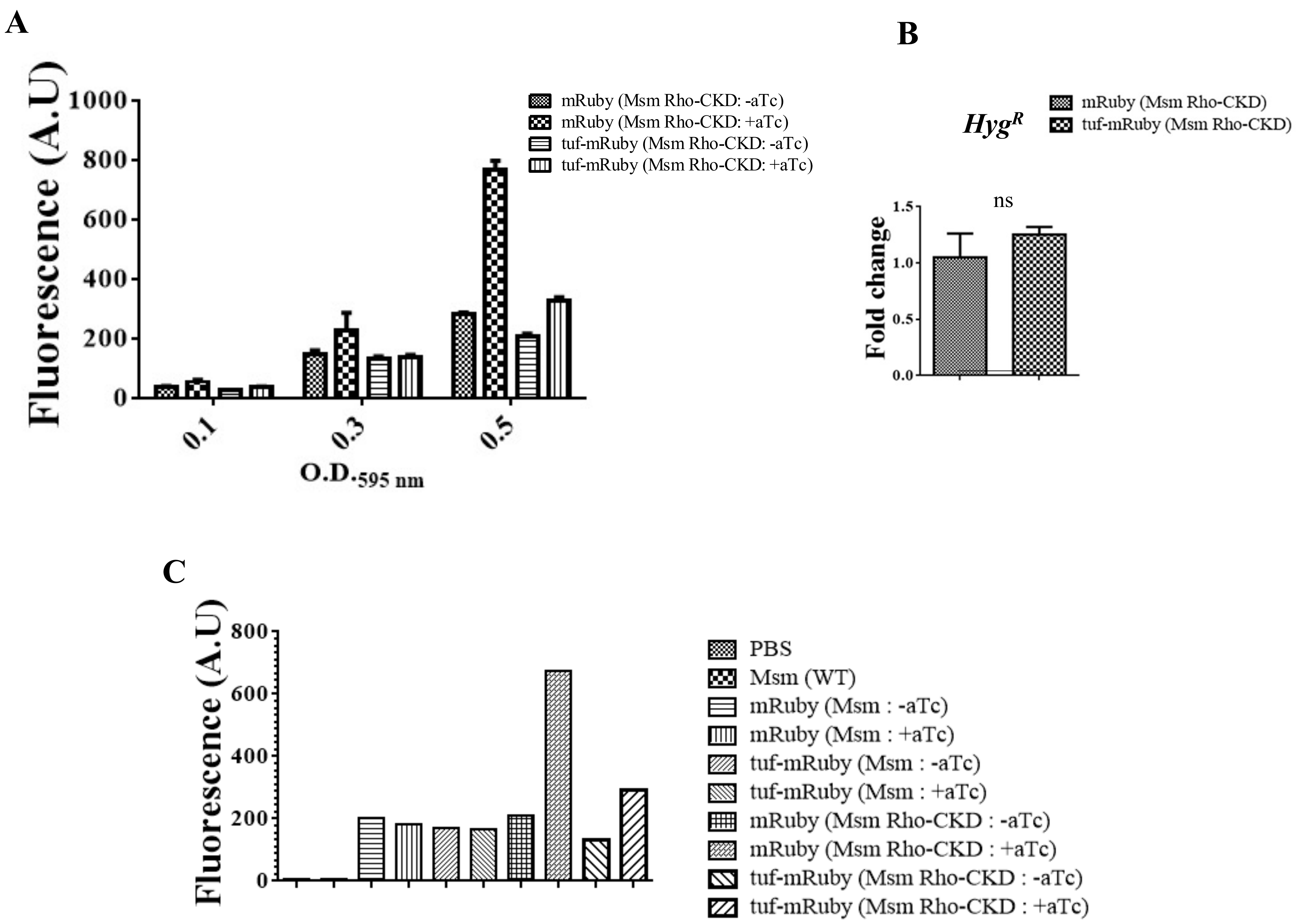

***Assumption: mRuby (Msm Rho CKD: +aTc) = 100% readthrough (No termination)***

$$\textit{Termination \% of tuf (Intrinsic terminator alone)} = \frac{\textit{((mRuby (Msm Rho CKD: +aTc) – (tuf-mRuby (MsmRho CKD: +aTc))}}{\textit{(mRuby (Msm Rho CKD: +aTc)}} \times 100$$

$$\textit{Termination \% of tuf (intrinsic terminator alone)} = \frac{675.5-293.5}{675.5} \times 100$$

***Termination % of tuf (Intrinsic terminator alone)= 56.55%***

$$\textit{Total termination \% of tuf (Intrinsic terminator alone + Rho alone)} = \frac{\textit{((mRuby (Msm Rho CKD: +aTc) – (tuf-mRuby (Msm Rho CKD: -aTc))}}{\textit{(mRuby (Msm Rho CKD: +aTc)}} \times 100$$

$$\textit{Total termination \% of tuf (Intrinsic terminator alone + Rho alone)} = \frac{675.5-130}{675.5} \times 100$$

***Termination % of tuf (Intrinstic termination + Rho)= 80.75%***

**Supplementary Figure 2. mRuby expression profile for *tuf* intrinsic terminator and estimation of the contribution intrinsic terminator and Rho in transcription termination (A)** mRuby expression profile for *tuf* intrinsic terminator at 0.1, 0.3 and 0.5 O.D$_{595\ nm}$ for Msm Rho-CKD strains in the presence and absence of aTc. **(B)** qRT-PCR to compare the levels of hygromycin resistance gene (*Hyg*$^{R}$) between Msm Rho-CKD strains described in figure A at 0.5 O.D$_{595\ nm}$ .**(C)** The difference in the mRuby expression in the presence and absence of intrinsic terminator at normal and reduced level of Rho was employed to estimate the individual contribution from both the modes of termination.

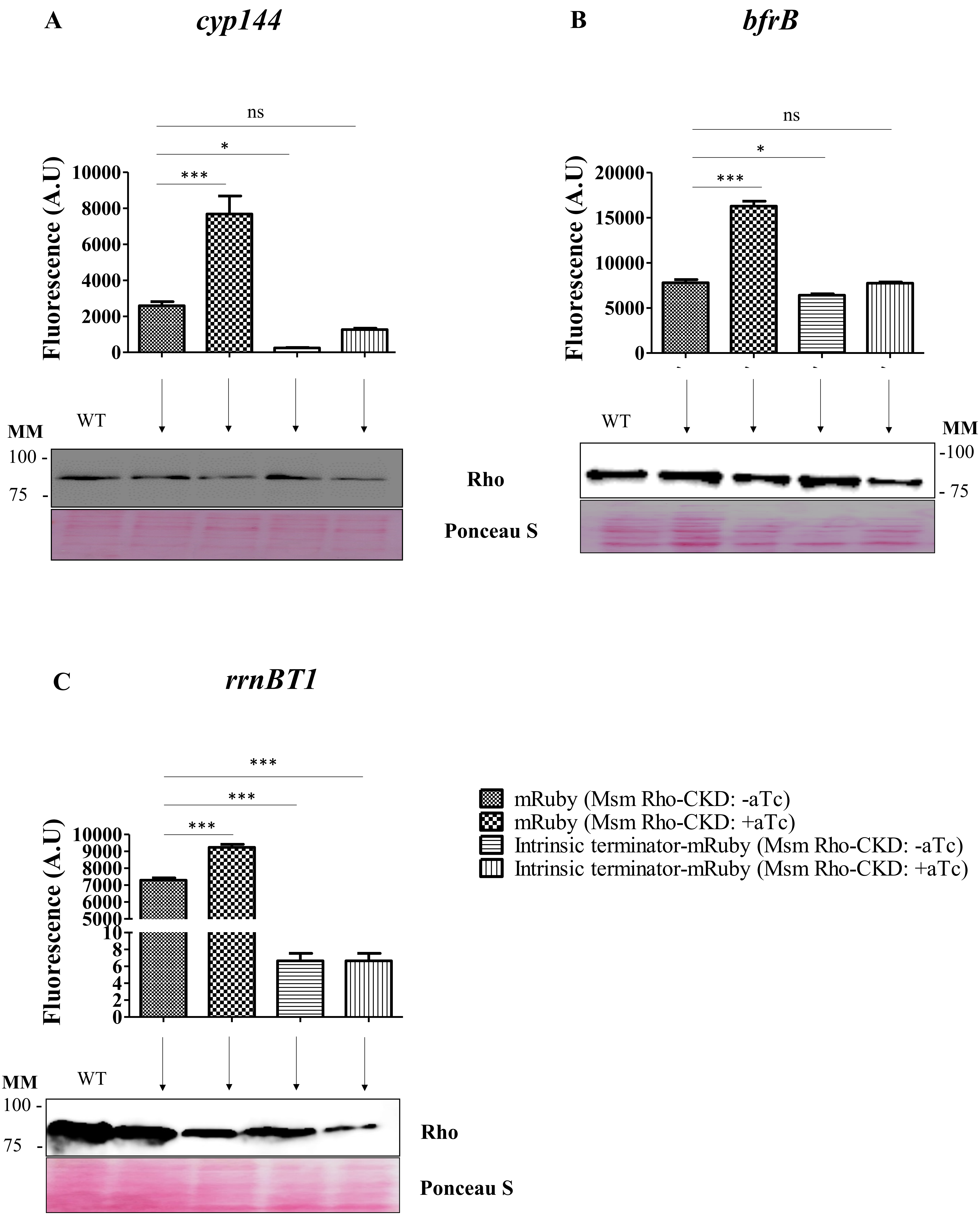

**Supplementary Figure 3. mRuby expression profile to estimate the contribution intrinsic terminator and Rho in transcription termination (A, B and C)** mRuby expression level for *cyp144, bfrB* and *rrnBT1* intrinsic terminators between Msm Rho-CKD strain in the presence and absence of aTc. Immunoblots showing Rho levels in these conditions and Ponceau S staining is shown as a loading control. The data represented is mean ± SD from the three independent experiments. The P-values were estimated using one-way ANOVA followed by Tukey multiple-comparisons test, *p<0.05, **p<0.001,***p<0.0001, ns – not significant.

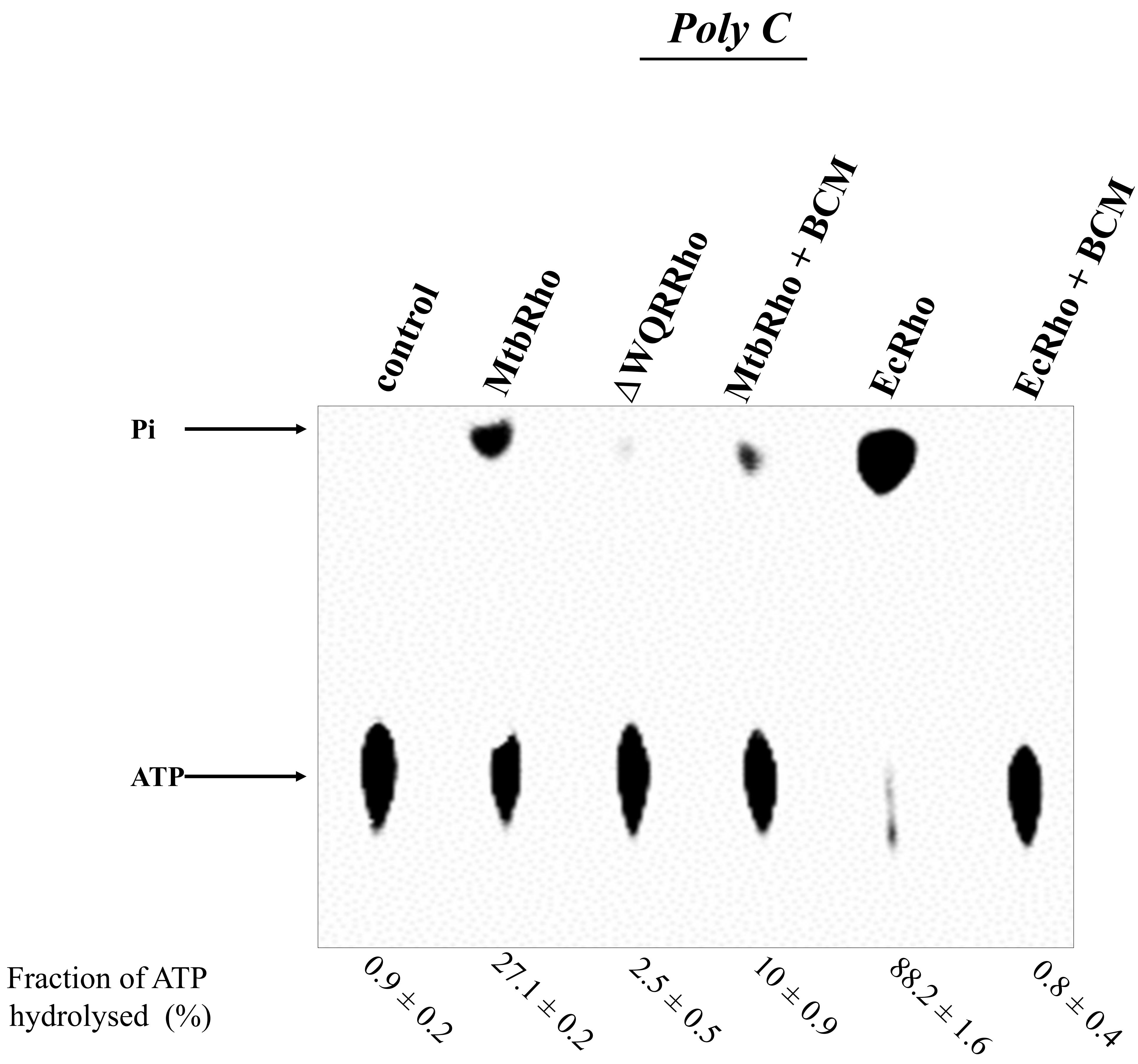

**Supplementary Figure 4. ATPase activity of MtbRho in the presence of Poly C.** ATPase assay was carried out using Poly C (40 ng) substrate in the presence of MtbRho, ΔWQRRho and EcRho (100 nM) . Rho-dependent ATP hydrolyses was inhibited in the presence of BCM (600 μM). The released inorganic phosphate (Pi) was visualized using Phosphorimager. Pi release was calculated with SDs determined from three independent experiments.

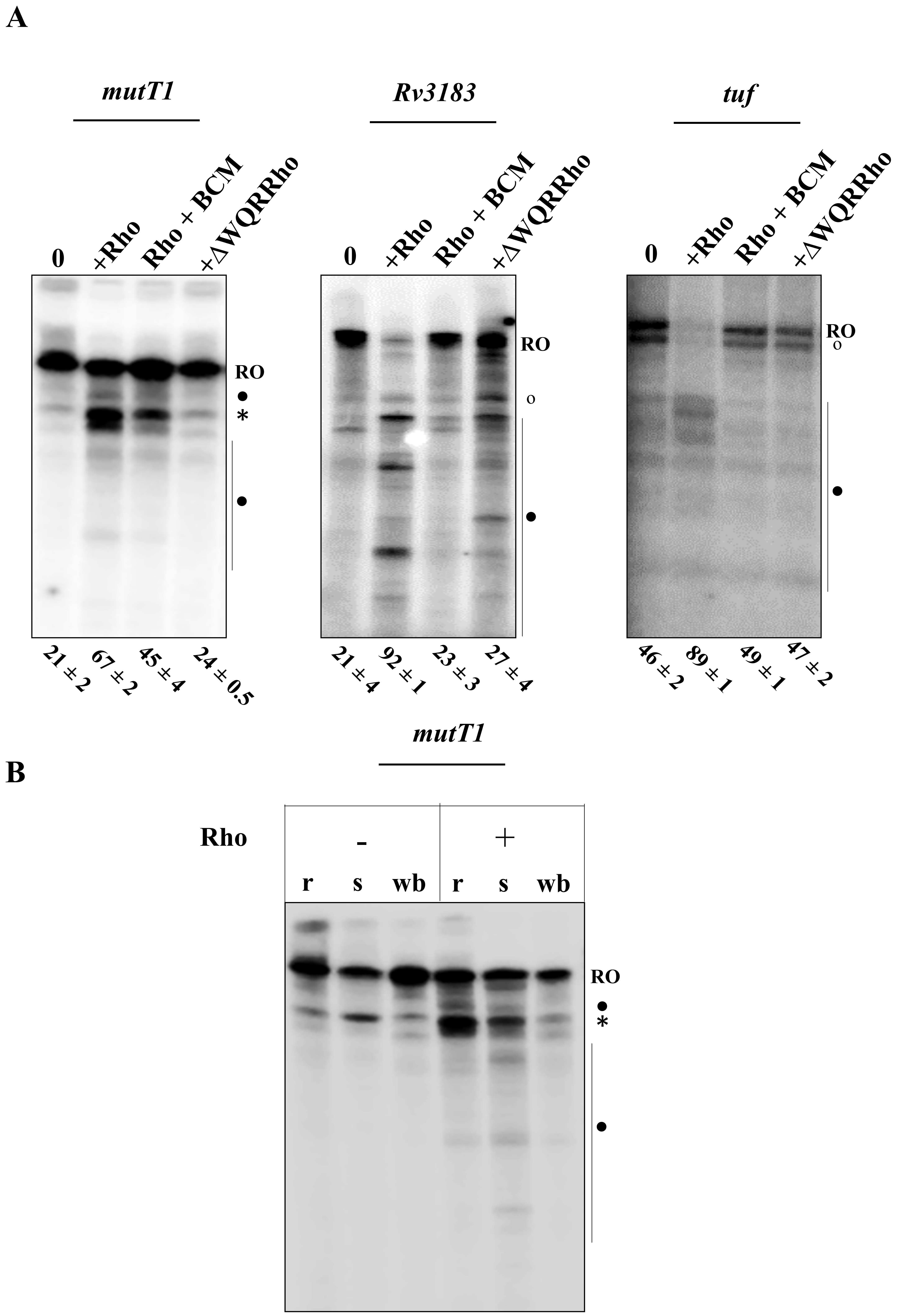

**Supplementary Figure 5. Rho control of transcriptional read-through from intrinsic terminators (A)** *In vitro* transcription assays were carried out using 10 nM linear DNA templates containing *mutT1, Rv3183,* and *tuf* intrinsic terminators in the presence and absence of MtbRho (100 nM), ΔWQRRho (100 nM) and BCM (600 μM). **(B)** Biotinylated linear DNA templates containing *mutT1* intrinsic terminator immobilized on streptavidin beads was subjected to *in vitro* transcription assay as described in Materials and methods, in the presence and absence of MtbRho (100 nM) to detect release of termination products. Transcription reaction mixtures were separated into supernatant (s) and washed-beads (wb) fractions for comparison to total reaction product (r). For both the assays transcripts were resolved on 6% 8M urea-PAGE. RO represents the run-off transcripts. Intrinsic termination (○), Rho-dependent termination (•) and Rho-IT termination products (*) were marked.

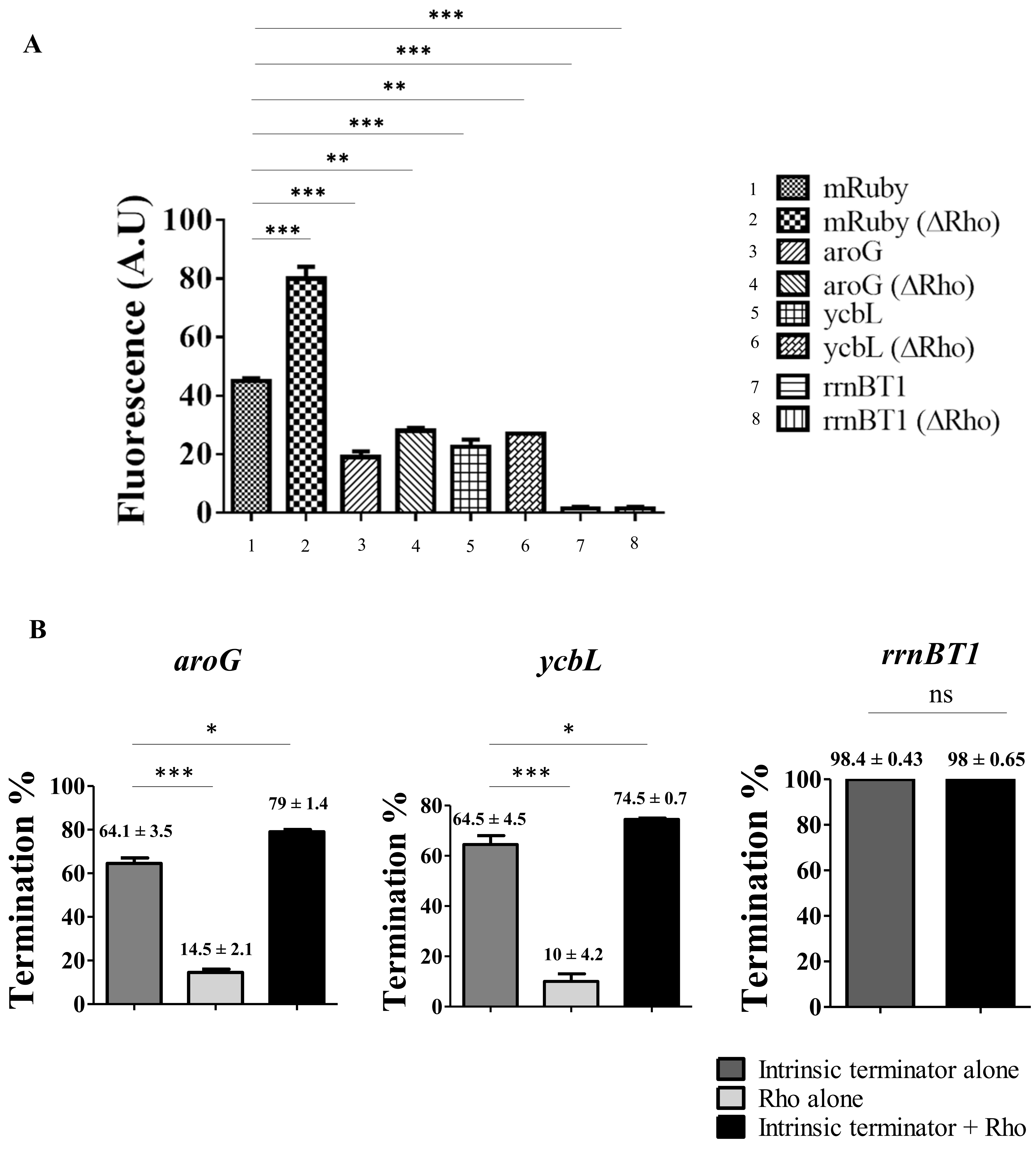

**Supplementary Figure 6. Evaluation of intrinsic terminator and Rho participation in *E. coli* (A)** mRuby expression levels with intrinsic terminators of *aroG, ycbL,* and *rrnBT1* in the presence and absence of Rho, in *E. coli RS1309* **(B)** Estimation of contribution of intrinsic terminator and Rho, both together and alone on the transcription termination efficiency of *aroG, ycbL,* and *rrnBT1* genes. The data represented is mean ± SD from the three independent experiments. The P-values were calculated by one-way ANOVA followed by Tukey multiple-comparisons test, *p<0.05, **p<0.001,***p<0.0001, ns – not significant.

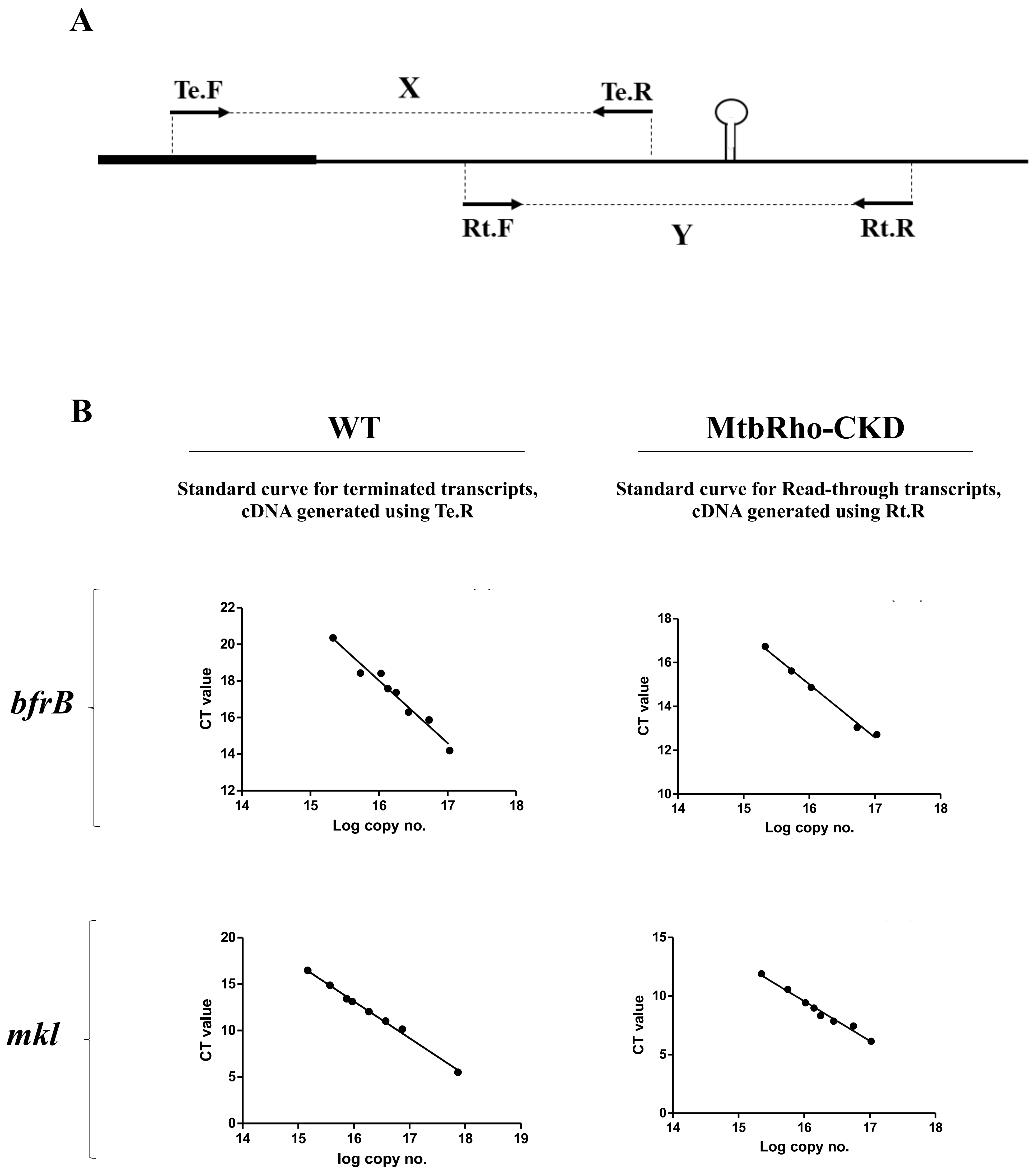

**Supplementary Figure 7. Estimation of terminated and read-through transcripts by RT-qPCR. (A)** Schematic of the RT-qPCR experimental design. Te.F-Te.R and Rt.F-Rt.R represent the primer sets to measure terminated and read-through transcripts respectively as described in Materials and methods. **(B)** Standard curves were generated using the known concentrations for *bfrB* and *mkl* transcripts to interpolate the levels of terminated and read-through transcripts in wild-type (WT) and MtbRho-CKD strain (Figure 3D) .

| | Gene | Organism | Gene sequence cloned | G+C content (%) | GeSTer predicted terminator sequence | *ΔG* (kcal/mol) | in vitro transcription with linear DNA template | | Remark | Reference |
|---|---|---|---|---|---|---|---|---|---|---|
| | | | | | | | run-off length | intrinsic termination product length | | |
| 1 | mutT1 | MtbH37Rv | TGTGCACCCCGACAAGTCCCGCAATCGCAAAGGCAGCACGTGGGTGTTGTCGTTGTCAGCCGGCAGGCTTGTGACAGCCGACCACATCGGCGGTGCGCTGGCCGCCAACGTGCGGGCCTAACACACGGATACCCTTCGTCACATTGCCACCGTGCAAAGGGTATCCGTGTGTCTTGACCTATTTGCGACCCCGCCGAGCGGTTGCCTTCTTGGCGGGAGCCTTGGTAGCCGGCCGCTTGGCCGCTGCCTTCTTTGCCGGCGCCTTGGTCGCCGCCTTACGCACCGATGCCTTGACCGCGGT | 63% | GATACCCTTCGTCACATTGCCACCGTGCAAAGGGTATCCGTGTGTCTTGACCTATTT | -15.68 | 362 nts | 219 nts | 1) mutT1 sequence cloned in pUC18-T7A1 plasmid downstream of T7A1 promoter to generated linear DNA template for in vitro transcription assays.<br>2) pUC18-T7A1-mutT1 was used to generate RNA for ATPase assays by using T7 RNA polymerase. | Ahmad et al., 2019 |
| 2 | Rv3183 | MtbH37Rv | CACCCTGCAGGCCTACGTTGCCGCCCTGGGCGGGCACCTGCGCATCGTCGCTGAGTTCGGCGAAAATACTGTCGAGCTGACCGCCTGAGCTAACTCACGCCCACACTTCCGGCCGGTCTCGATCTCCCAAGCCCCAGCACAGCTCGTGTTCCCAATCTGTTCCCAACCAGATCCTTAGCTATGCGCATGTTCCCAAAAGTGTTCCCGCCCATGAAAACGGCCCCCGGAGTCTCCTCCGAGGGCCATTTCGCCGGTAGCGGGGACAGGATTCGAACCTGCGACCTCTGGGTTATGAGCTAACCAGTCGCAATCTCTCCCATCGCGGTCGGTCTCATACGTCCAGATCAGCCTCTATTCCGCCGTCCAGCCTGTTCCGCCGCGTCGCGGTTGTACGGATTTGAACCGCCCCGGCATGTCCGGAGACTCCAGTTCTTGGAAAGGATGGGGTCATGTCAGGTGGTTCATCGAGGAGGTACCCGCCGGAGCTGCGTGAGCGGGCGGTGCGGATGGTCG | 67% | GGCCCCCGGAGTCTCCTCCGAGGGCCATTTCGCCGG | -20.3 | 574 nts | 289 nts | 1) Rv3183 sequence cloned in pUC18-T7A1 plasmid downstream of T7A1 promoter to generated linear DNA template for in vitro transcription assays.<br>2) pUC18-T7A1-Rv3183 was used to generate RNA for ATPase assays by using T7 RNA polymerase. | Ahmad et al., 2019 |
| 3 | mkl | MtbH37Rv | TGTTCGGGAGCTGATCACGCAACGTTTGTATCTGCTGCCGACCTTCCGTTGGCGGCTCGCGTAGGTGGCACAGTCCGCGAAGTGCTTGGGCCGCTGATCAAGGCGCTCCCGGAGCACAATCCAGACATGTCAGGCCGTCACCGACGCACAGGCGACGGCCCTCGAGCAGCGTGGGAAGAGCCGGGCTCGTCGAGTGGATCACACATTTCGAGGCGCTCTCGTGTATCGAGCGGCACATCAGCCATGCGTGTCTCCTTGTCCTGCCTTCTCCAGAGGAA | 61% | GGCCGTCACCGACGCACAGGCGACGGCCCTCGAGCAGC | -17.15 | 339 nts | 205 nts | 1) mkl sequence cloned in pUC18-T7A1 plasmid downstream of T7A1 promoter to generated linear DNA template for in vitro transcription assays.<br>2) pUC18-T7A1-mkl was used to generate RNA for ATPase assays by using T7 RNA polymerase. | Ahmad et al., 2019 |
| 4 | bfrB | MtbH37Rv | GGATCAGCGAGTGGTCCCGTTCGCCCGCCCGTCTTCCAGCCAGGCCTTGGTGCGGCCGGGGTGGTGAGTACCAATCCAGGCCACCCCGACCTCCCGGCAAAAGTCGATGTCCTCGTACTCATCGACGTTCCAGCAGTACACCGCCCGGCCCTGAGCTGCCGAGCGGTCAACGAGTTGCGGATATTCCTTTAACGCAGGCAGTGAGGGTCCCACGGCGGTTGCCCCGACCGCCGTGGCCGCACTGCTGGTCAGGTATCGGGGGGTCTTGCCGAGCAACACCGTCGGCA | 65% | CCACGGCGGTTGCCCCGACCGCCGTGGCCGCACTGCT | -20.09 | 348 nts | 281 nts | 1) bfrB sequence cloned in pUC18-T7A1 plasmid downstream of T7A1 promoter to generated linear DNA template for in vitro transcription assays.<br>2) pUC18-T7A1-bfrB was used to generate RNA for ATPase assays by using T7 RNA polymerase.<br>3) bfrB sequence cloned downstream of Hsp60 promoter in pMV261-mRuby plasmid for mRuby reporter fluorescence assays. | Ahmad et al., 2019 |
| 5 | metE | MtbH37Rv | GGTGCTCGACGACCTGAACGCGATCGGCTTCGCCAACGGTGTGGGCCCGGGCGTCTATGACATTCACTCGCCACGGGTGCCCTCCGCTGAGGAGATGGCCGACTCGTTGCGGGCCGCGTTGCGCGCGGTGCCGGCCGAGCGGCTGTGGGTCAACCCCGACTGCGGACTGAAGACCCGCAATGTCGACGAGGTGACCGCGTCGCTGCACAACATGGTCGCCGCCGCCCGGGAGGTGCGCGCGGGCTAGCTGACGAGCAGCTAGCCCGGCAGCACGTGGACCGGCACGTCGTCGTCCCAGGGCTGCCGGTTCACCATGTCGGCCACCCGGACATAGCCGCGCTCGAACTCCCCGGTCGCCGCGTCTACCAACCGGTACTCCTGGGTTTGCCGATGGATGGGCTGCGCGTCGAGCAG | 69% | CGGGCTAGCTGACGAGCAGCTAGCCCGGCAGCACGTG | -22.45 | 475 nts | 311 nts | 1) metE sequence cloned in pUC18-T7A1 plasmid downstream of T7A1 promoter to generated linear DNA template for in vitro transcription assays.<br>2) pUC18-T7A1-metE was used to generate RNA for ATPase assays by using T7 RNA polymerase. | Ahmad et al., 2019 |
| 6 | cyp144 | MtbH37Rv | GCTGCACTGGCACGCTTGGAGGCTCGAATCGTCTTGCGTCTGCTGCTCGATCGCACCTCGGTAATTGAGGCAGCCGATGTCGGCGGGTGGTTGCCCAGTATCCTGGTGCGCCGCATCGAGCGGCTAGAGCTAGCTGTACAATAGGCGCTCGACGACTCCTATTGCAGCACAACGGATATCAGCAACAGCAGGTGCCAACCGCGGCGATCGGATGCGTGAGAATAGTGAAAGTGGTTGTCGCGGTCAGGATTTCTGCGATCAACCCTACCCGCATGACGCCGGCGGGTGGCCCCCGCCGGCCACGATAAATGCTTCGACCGCCGTGGCCCGCTCGTAACCTTCGACCTCCACGCGCCACTC | 61% | GCCGGCGGGTGGCCCCCGCCGGCCACGATAAAT | -20 | 421 nts | 345 nts | 1) cyp144 sequence cloned in pUC18-T7A1 plasmid downstream of T7A1 promoter to generated linear DNA template for in vitro transcription assays.<br>2) pUC18-T7A1-cyp144 was used to generate RNA for ATPase assays by using T7 RNA polymerase.<br>3) cyp144 sequence cloned downstream of Hsp60 promoter in pMV261-mRuby plasmid for mRuby reporter fluorescence assays. | Ahmad et al., 2019 |
| 7 | tuf | MtbH37Rv | CGGTGAAGTTGATCCAGCCCGTCGCCATGGACGAAGGTCTGCGTTTCGCGATCCGCGAGGGTGGCCGCACCGTGGGCGCCGGCCGGGTCACCAAGATCATCAAGTAGGTCTACCGGCCACCAGACGCAAAAGAACATGATGGGCGCACCAGCGCCCATCATGTTCTTTTGCGTCTGCTCGCGAAAATGCCCAGCGTGCGGCGCTACGCTGACATGGACCCTCCGACGAGGCAAGGAGCAGGCACGTGTTAGCGCGCTACATCAAGATGCAGTTATTGGTGCTGTTGTGCGGTGGTCTGGTCGGGCCGATCTTCTTGGTCGTCTACTTCACGCTCGGACTGGGCAGCCTGATGTCGTGGATGTTCTATGTCGG | 60% | CATGATGGGCGCACCAGCGCCCATCATGTTCTTTT | -23.22 | 433 nts | 221 nts | 1) tuf sequence cloned in pUC18-T7A1 plasmid downstream of T7A1 promoter to generated linear DNA template for in vitro transcription assays.<br>2) pUC18-T7A1-tuf was used to generate RNA for ATPase assays by using T7 RNA polymerase.<br>3) tuf sequence cloned downstream of Hsp60 promoter in pMV261-mRuby plasmid for mRuby reporter fluorescence assays. | This work |
| 8 | aroG | E. coli K-12 MG1655 | TACCGATGCTCTGTTACGTCAACTGGCGAATGCAGTAAAAGCGCGTCGCGGGTAAGGTTTAATTGTCGGATGCGCCGTCAGAGTGGCGTATCCGATGAATCACCACAGGCCTGATAAGTCGCGCAGCGTCGCATCAGGCAATGTGCTCCATTGTTAGCAACAAAAAAGCCGACTCACTTGCAGTCGGCTTTCTCATTTTAAACGAATGACGTTTACTTCGCTTTACCCTGGTTTGCAACCGCCGCTGCTTTCGCTGCGATCTCGTCAGCATTACCCAGATAATAGCGTTTCAGCGGTTTGAAATTCTCGTCGAACTCATACACCA | 49% | AAAAGCCGACTCACTTGCAGTCGGCTTTCTCATTTTAAA | -12.52 | 386 nts | 234 nts | 1) aroG sequence cloned in pUC18-T7A1 plasmid downstream of T7A1 promoter to generated linear DNA template for in vitro transcription assays.<br>2) aroG sequence cloned downstream of Hsp60 promoter in pMV261-mRuby plasmid for mRuby reporter fluorescence assays. | This work |
| 9 | ycbL | E. coli K-12 MG1655 | TGCCCGTCTGGTAAGGCACATAAAAAAGCCCGCTTTTAATGCTGGCCTGGATTTCTGGCAAAGTGCGCTTTGTTTATGCCGGATGCGGCACGAGCGCCTTATCCGGCCTACAAAATCGTGCAAATTCAAAATATTGCAGGGGACGCGTAGGCCTGATAAGCGTAGCGCATCAGGCAATGTTGCGTTTGTCATCAGTCTCAGCCCGCTTTTCAGCGGGCTTCATTGTTTTTAATGCTTACAGCACTGCCACAATCGC | 50% | AGCCCGCTTTTCAGCGGGCTTCATTGTTTTT | -13.98 | 345 nts | 265 nts | 1) ycbL sequence cloned in pUC18-T7A1 plasmid downstream of T7A1 promoter to generated linear DNA template for in vitro transcription assays.<br>2) ycbL sequence cloned downstream of Hsp60 promoter in pMV261-mRuby plasmid for mRuby reporter fluorescence assays. | This work |
| 10 | rrnBT1 | E. coli K-12 MG1655 | GTCCCACCTGACCCCATGCCGAACTCAGAAGTGAAACGCCGTAGCGCCGATGGTAGTGTGGGGTCTCCCCATGCGAGAGTAGGGAACTGCCAGGCATCAATAAAACGAAAGGCTCAGTCGAAAGACTGGGCCTTTCGTTTTATCTGTTGTTTGTCGGTGAACGCTCTCCTGAGTAGGACAAATCCGCCGGGAGCGGATTTGAACGTTGCGAAGCAACGGCCCGGAGGGTGGCGGGCAGG | 57% | GGCTCAGTCGAAAGACTGGGCCTTTCGTTTT | -17 | 300 nts | 176 nts | 1) rrnBT1 sequence cloned in pUC18-T7A1 plasmid downstream of T7A1 promoter to generated linear DNA template for in vitro transcription assays.<br>2) pUC18-T7A1-rrnBT1 was used to generate RNA for ATPase assays by using T7 RNA polymerase.<br>3) rrnBT1 sequence cloned downstream of Hsp60 promoter in pMV261-mRuby plasmid for mRuby reporter fluorescence assays. | This work |

**Supplementary Table 1.** Sequence and other details of intrinsic terminators used in the study.

| Organism | Gene | Termination efficiency | Reference |
|---|---|---|---|
| Mycobacterium tuberculosis | *Rv3444c* | 56 ± 7 | czyz et al, 2014 |
| | *tuf* | 57 ± 1 | |
| | *Rv1324* | 15 ± 2 | |
| | *rpoC* | 14 | |
| | *mutT1* | 21 ± 2 | Ahmad et al, 2019 |
| | *Rv3183* | 21 ± 4 | |
| | *Lsr2* | 15 ± 2 | |
| | *mkl* | 55 ± 3 | |
| | *metE* | 33 ± 3 | |
| | *cyp144* | 70 ± 2 | |
| | *bfrB* | 42 ± 1 | |
| Escherichia coli | *trpA* | 80 ± 4 | czyz et al, 2014 |
| | *rpoC* | 80 | |
| | *rrnBT1* | 75 ± 6 | |
| | *galM* | ND | Wang et al, 2018 |
| Bacillus subtilis | *yetJ** | 47 | Mandell et al, 2021 |
| | *ktrD** | 66 | |
| | *metS* | 70 | Mondal et al, 2016 |
| | *ypmT** | 23 | |
| | *yqxC** | 33 | |
| | *yusV* | 60 | |
| | *argI** | 46 | |
| | *bioyB** | 31 | |
| | *proS** | 68 | |
| | *trpL* | 79 | |
| | *thrS* | 18 | |
| | *yvbJ* | 38 | |
| | *yetH** | 35 | |
| | *ilvD* | 33 | |
| Phage lambda | *λtI* | 5 0 | |
| | *λtr2* | 67 | |
| T7 | *TTΦ-U6G(cu)* | 73 ± 2 | Molodtsov et al., 2014 |
| SP6 | *SP6 terminator* | 45±2 | Yoo and Kang, 2000 |
| actinophage ΦC31 | *et* [a] | 75 | Ingham CJ et al, 1995 |
| | *et* [b] | 41 | |
| | *et* [c] | 67 | |
| Streptococcal plasmid pMV158 | *TII* | 87.5 | Gloria del Solar1 and Manuel Espinosa, 2001 |
| | *T7Te* | 88 | Reynolds et al., 1991 |
| | *RNA I* | 73 | |
| | *BS6* | 30 | |
| | *BS7* | 28 | |
| | *P14* | 25 | |
| | *tonB* | 19 | |
| | *T3Te* | 14 | |

**Supplementary Table 2. Known intrinsic terminators and their termination efficiency.** List of few known intrinsic terminators with their termination efficiency *in vitro* . * represents the termination efficiency in the presence of Nus factor. ND is Not determined.

**Bacterial Strains**

| | Strain name/no. | Description | Reference |
|---|---|---|---|
| 1 | MtbRho-CKD | MtbH37Rv, Carrying pRH2520 and pRH2521 expressing dcas9 and MtbRho sgRNA respectively | This work |
| 2 | MsmRho-CKD | Msm mc2155, Carrying pRH2520 and pRH2521 expressing dcas9 and MsmRho sgRNA respectively | This work |
| 5 | Msm SM07 | An isogenic strain to Msm mc2155 having the replacement of rpoC with a His-tagged rpoC | Mukherjee et al, 2008 |
| 3 | AMO14 | E. coli strain with chromosomal rho gene inactivated, rho supplied in trans on plasmid with temperature-sensitive origin of replication | Martinez et al ., 1996 |
| 4 | RS1309 | E. coli MG1655 Δrho Δrac, Carrying shelter plasmid pHYD1201 (AmpR) | Valabhoju et al., 2016 |
| 6 | E. coli BL21(DE3) | (hsdS gal (lcIts857 ind1 Sam7 nin5 lacUV5-T7 gene 1) used for expression and purification | Laboratory stock |
| 7 | E. coli DH10B | Δ(mrr-hsd RMS-mcrBC) mcrA recA1) used for cloning | Laboratory stock |

**Plasmid used in the study**

| | Plasmid names | Description | Reference |
|---|---|---|---|
| 1 | pRH2520 | Integrative plasmid carrying an inactive version of Streptococcus pyogenes cas9, $Kan^R$ | Singh et al., 2016 |
| 2 | pRH2521 | Plasmid for cloning gene specific sgRNA downstream of TetR-regulated smyc promoter (Pmyc1tetO), $Hyg^R$ | Singh et al., 2016 |
| 3 | pRH2521-MtbRho-CKD | pRH2521 plasmid carrying MtbRho specific sgRNA, $Hyg^R$ | This work |
| 4 | pRH2521-MsmRho-CKD | pRH2521 plasmid carrying MsmRho specific sgRNA, $Hyg^R$ | This work |
| 5 | pMV261-mRuby | mRuby cloned downstream of hsp60 promoter at EcoRI/HindIII site of pMV261, $Kan^R$ | This work |
| 6 | pMV261-tuf-mRuby | DNA segment carrying stop-to-stem region of tuf cloned upstream of mRuby at BamHI/EcoRI site of pMV261-mRuby, $Kan^R$ | This work |
| 7 | pMV261-bfrB-mRuby | DNA segment carrying stop-to-stem region of bfrB cloned upstream of mRuby at PvuII/EcoRI site of pMV261-mRuby, $Kan^R$ | This work |
| 8 | pMV261-cyp144-mRuby | DNA segment carrying stop-to-stem region of cyp144 cloned upstream of mRuby at BamHI/EcoRI site of pMV261-mRuby, $Kan^R$ | This work |
| 9 | pMV261-rrnBT1-mRuby | DNA segment carrying stop-to-stem region of rrnBT1 cloned upstream of mRuby at BamHI/EcoRI site of pMV261-mRuby, $Kan^R$ | This work |
| 10 | pMV261-aroG-mRuby | DNA segment carrying stop-to-stem region of aroG cloned upstream of mRuby at BamHI/PstI site of pMV261-mRuby, $Kan^R$ | This work |
| 11 | pMV261-ycbL-mRuby | DNA segment carrying stop-to-stem region of ycbl cloned upstream of mRuby at BamHI/PstI site of pMV261-mRuby, $Kan^R$ | This work |
| 12 | pRH2521-MsmRho-CKD-mRuby | DNA segment from Hsp60 promoter to mRuby subcloned from pMV261-mRuby at NotI/HpaI site of pRH2521-MsmRho-CKD , $Hyg^R$ | This work |
| 13 | pRH2521-MsmRho-CKD-tuf-mRuby | DNA segment from Hsp60 promoter to mRuby subcloned from pMV261-tuf-mRuby at NotI/HpaI site of pRH2521-MsmRho-CKD , $Hyg^R$ | This work |
| 14 | pRH2521-MsmRho-CKD-bfrB-mRuby | DNA segment from Hsp60 promoter to mRuby subcloned from pMV261-bfrB-mRuby at NotI/HpaI site of pRH2521-MsmRho-CKD , $Hyg^R$ | This work |
| 15 | pRH2521-MsmRho-CKD-cyp144-mRuby | DNA segment from Hsp60 promoter to mRuby subcloned from pMV261-cyp144-mRuby at NotI/HpaI site of pRH2521-MsmRho-CKD , $Hyg^R$ | This work |
| 16 | pRH2521-MsmRho-CKD-rrnBT1-mRuby | DNA segment from Hsp60 promoter to mRuby subcloned from pMV261-rrnBT1-mRuby at NotI/HpaI site of pRH2521-MsmRho-CKD , $Hyg^R$ | This work |
| 17 | pRH2521-$mRuby^E$ | DNA segment from Hsp60 promoter to mRuby subcloned from pMV261-mRuby at NotI/HpaI site of pRH2521, $Hyg^R$ | This work |
| 18 | pRH2521-aroG-$mRuby^E$ | DNA segment from Hsp60 promoter to mRuby subcloned from pMV261-aroG-mRuby at NotI/HpaI site of pRH2521, $Hyg^R$ | This work |
| 19 | pRH2521-ycbL-$mRuby^E$ | DNA segment from Hsp60 promoter to mRuby subcloned from pMV261-ycbL-mRuby at NotI/HpaI site of pRH2521, $Hyg^R$ | This work |
| 20 | pRH2521-rrnBT1-$mRuby^E$ | DNA segment from Hsp60 promoter to mRuby subcloned from pMV261-rrnBT1-mRuby at NotI/HpaI site of pRH2521, $Hyg^R$ | This work |
| 21 | pET11d-MtbRho | MtbRho cloned in NcoI and BamHI sites of pET11d, $Amp^R$ | Mitra et al., 2014 |
| 22 | pET11d-ΔWQRRho | Walker B motif, Q-loop, and R-loop of the ATPase domain of Rho deleted from pET11d-MtbRho, $Amp^R$ | This work |
| 23 | pET21a-EcRho | E. coli Rho with C-terminal hexahistidine tag, cloned in pET21a, $Amp^R$ | Epshtein et al, 2010 |
| 24 | pUC18-T7A1-mutT1 | DNA segment carrying stop-to-stem region of mutT1 cloned downstream of T7A1 promoter at XbaI/HindIII site of pUC18-T7A1, $Amp^R$ | Ahmad et al., 2019 |
| 25 | pUC18-T7A1-Rv3183 | DNA segment carrying stop-to-stem region of Rv3183 cloned downstream of T7A1 promoter at XbaI/HindIII site of pUC18-T7A1, $Amp^R$ | Ahmad et al., 2019 |
| 26 | pUC18-T7A1-mkl | DNA segment carrying stop-to-stem region of mkl cloned downstream of T7A1 promoter at XbaI/HindIII site of pUC18-T7A1, $Amp^R$ | Ahmad et al., 2019 |
| 27 | pUC18-T7A1-bfrB | DNA segment carrying stop-to-stem region of bfrB cloned downstream of T7A1 promoter at XbaI/HindIII site of pUC18-T7A1, $Amp^R$ | Ahmad et al., 2019 |
| 28 | pUC18-T7A1-metE | DNA segment carrying stop-to-stem region of metE cloned downstream of T7A1 promoter at XbaI/HindIII site of pUC18-T7A1, $Amp^R$ | Ahmad et al., 2019 |
| 29 | pUC18-T7A1-cyp144 | DNA segment carrying stop-to-stem region of cyp144 cloned downstream of T7A1 promoter at XbaI/HindIII site of pUC18-T7A1, $Amp^R$ | Ahmad et al., 2019 |
| 30 | pUC18-T7A1-aroG | DNA segment carrying stop-to-stem region of aroG cloned downstream of T7A1 promoter at XbaI/HindIII site of pUC18-T7A1, $Amp^R$ | This work |
| 31 | pUC18-T7A1-ycbL | DNA segment carrying stop-to-stem region of ycbL cloned downstream of T7A1 promoter at XbaI/HindIII site of pUC18-T7A1, $Amp^R$ | This work |

**Supplementary Table 3.** Strains and plasmids used in this study

| Primer | Sequence 5'-3' |
|---|---|
| mutT1Fwd | AGCTCTAGATGTGCACCCCGACAAG |
| mutT1Rev | AGCAAGCTTACCGCGGTCAAGGCATC |
| Rv3183 Fwd | CGATCTAGACACCCTGCAGGCCTACG |
| Rv3183 Rev | CGAAAGCTTCGACCATCCGCACCGC |
| mkl Fwd | CGATCTAGATGTTCGGGAGCTGATCACG |
| mkl Rev | CGAAAGCTTTTCCTCTGGAGAAGGCAGG |
| bfrB Fwd | CGATCTAGAGGATCAGCGAGTGGTCCCG |
| bfrB Rev | CGAAAGCTTTGCCGACGGTGTTGCTCG |
| metE Fwd | CGATCTAGAGGTGCTCGACGACCTGAACGCG |
| metE Rev | CGAAAGCTTCTGCTCGACGCGCAGCCC |
| cyp144 Fwd | CGATCTAGATGGTGCGCCGCATCGAGC |
| cyp144 Rev | CGAAAGCTTAGTGGCGCGTGGAGGTCG |
| tuf Fwd | CGATCTAGAGTCTACCGGCCACCAGA |
| tuf Rev | CGAAAGCTTCGGCCCGACCAGACCAC |
| aroG Fwd | CGCGGATCCTACCGATGCTCTGTTACGTC |
| aroG Rev | CGCCTGCAGTGGTGTATGAGTTCGACGAG |
| ycbL Fwd | CGCGGATCCTGCCCGTCTGGTAAGGCAC |
| ycbL Rev | CGCCTGCAGGCGATTGTGGCAGTGCTGTAAGC |
| rrnBT1 Fwd | CGATCTAGAGTCCCACCTGACCCCA |
| rrnBT1 Rev | CGAAAGCTTCCTGCCCGCCACCC |
| mRuby Fwd | CGATCTAGACGGTGACCACAACGCGCC |
| mRuby Rev | CGATCTAGATTACTTGTACAGCTCGTCCATCCCACCAC |
| mRuby RT Fwd | GTCGTTCATGTATGGCAGCC |
| mRuby RT Rev | TCACGGGACCATTGGAGG |
| MtbRho RT Fwd | CCCAAAGCGGGCAAGAC |
| MtbRho RT Rev | TGGCATTCCGGGTTGTTC |
| bfrB-RT-RO Fwd | GGATCAGCGAGTGGTCCCG |
| bfrB-RT-RO Rev | CTCAGGGCCGGGCGGTGTAC |
| bfrB-RT-T Fwd | GTACACCGCCCGGCCCTGAG |
| bfrB-RT-T Rev | TGCCGACGGTGTTGCTCG |
| mkl-RT-RO Fwd | TGTTCGGGAGCTGATCACG |
| mkl-RT-RO Rev | GCCTGACATGTCTGGATTGTGC |
| mkl-RT-T Fwd | GCACAATCCAGACATGTCAGGC |
| mkl-RT-T Rev | CCTCTGGAGAAGGCAGGACAAG |
| Hyg-RT-Fwd | GCTCACCACCCATTCCGAG |
| Hyg-RT-Rev | CAGCAGCGTGTCCACGTC |

**Supplementary Table 4.** Sequences of primers used in this study
